# Single cell preparations of *Mycobacterium tuberculosis* damage the mycobacterial envelope and disrupt macrophage interactions

**DOI:** 10.1101/2022.06.16.496372

**Authors:** Ekansh Mittal, Andrew T. Roth, Anushree Seth, Srikanth Singamaneni, Wandy Beatty, Jennifer A. Philips

## Abstract

For decades, investigators have studied the interaction of *Mycobacterium tuberculosis* (Mtb) with macrophages, which serve as a major cellular niche for the bacilli. Because Mtb are prone to aggregation, investigators rely on varied methods to disaggregate the bacteria for these studies. Here, we examined the impact of routinely used preparation methods on bacterial cell envelop integrity, macrophage inflammatory responses, and intracellular Mtb survival. We found that both gentle sonication and filtering damaged the mycobacterial cell envelope and markedly impacted the outcome of macrophage infections. Unexpectedly, sonicated bacilli were hyperinflammatory, eliciting dramatically higher TLR2-dependent gene expression and elevated secretion of IL-1β and TNF-α. Despite evoking enhanced inflammatory responses, sonicated bacilli replicated normally in macrophages. In contrast, Mtb that had been passed through a filter induced little inflammatory response, and they were attenuated in macrophages. Previous work suggests that the mycobacterial cell envelope lipid, phthiocerol dimycocerosate (PDIM), dampens macrophage inflammatory responses to Mtb. However, we found that the impact of PDIM depended on the method used to prepare Mtb. In conclusion, widely used methodologies to disaggregate Mtb may introduce experimental artifacts in Mtb-host interaction studies, including alteration of host inflammatory signaling, intracellular bacterial survival, and interpretation of bacterial mutants.

## Introduction

A fundamental feature of the pathogenesis of *Mycobacterium tuberculosis* (Mtb), the etiologic agent of tuberculosis (TB), is its ability to survive and grow in host macrophages. For more than five decades, many laboratories have investigated how Mtb interacts with and modulates the function of macrophages. Mtb is characterized by a “waxy” coat, which confers its distinctive acid-fast staining properties. The complex cell envelope is important for pathogenesis and also allows Mtb to withstand adverse conditions (Dulberger et al., 2020). The mycobacterial envelope consists of a plasma membrane, peptidoglycan-arabinogalactan layer, outer membrane, and capsular layer (Dulberger et al., 2020). The outermost layers of the envelope are crucial in host-pathogen interactions given that they are directly able to interact with host cells. The capsule is composed primarily of a loose matrix of neutral polysaccharides (Kalscheuer et al., 2019), while the outer membrane is composed of long-chain mycolic fatty acids that are free, attached to trehalose, or covalently attached to the underlying arabinogalactan-peptidoglycan layer. The outer membrane also contains a complex array of unique lipids. Many of these lipids are bioactive; they can intercalate into host membranes, alter inflammatory signaling, disrupt phagosome maturation, and promote mycobacterial virulence (Cambier et al., 2020; Lerner et al., 2018; Quigley et al., 2017). Some outer membrane lipids, including phthiocerol dimycocerosate (PDIM), phenolic glycolipids, and sulfoglycolipids, are thought to act as antagonists of pathogen recognition receptors (PRRs) or to shield underlying pathogen associated molecular patterns (PAMPs) to prevent them from activating PRRs (Blanc et al., 2017; Cambier et al., 2014; Reed et al., 2004). Thus, the integrity of the envelope is crucial for host interactions and bacterial virulence.

Given the importance of Mtb-macrophage interactions, a mainstay of the experimental approach of many laboratories is the use of *in vitro* cultured Mtb to infect myeloid cells. However, the tendency of Mtb to form bacterial clumps has long presented an obstacle to these experiments, which depend on using precise and reproducible amounts of bacteria (Wells, 1946). For this reason, low concentrations of detergents are commonly added to culture media, but this does not fully resolve the problem.

Therefore, additional measures are routinely taken to generate single cell suspensions, including sonicating, syringing, centrifuging, filtering, vortexing with glass beads, or some combination of these procedures. The use of these techniques varies widely across different laboratories, the methodology used is not always reported, and there has been little consideration as to how these techniques impact experimental outcomes.

We are interested in how mycobacterial protein and lipid effectors modulate macrophage responses. Sometimes our results differed from published data, leading us to question whether the method of preparing the bacilli explained the differences. However, there was minimal literature into how dispersing mycobacterial clumps impacts the envelope and host-pathogen interactions. Previous studies demonstrated that use of detergent and agitation can release capsular constituents (Lemassu et al., 1996; Sani et al., 2010). In addition, it was reported that Mtb that had been sonicated for 90 seconds were better able to bind to macrophages, and the bacterial envelope appeared uneven and bulging on transmission electron microscopy (Stokes et al., 2004). Another study showed that passing bacterial cultures through 5μm pore filters improved reproducibility of high-throughput antibacterial drug screening compared to vortexing, but the impact on host-pathogen interactions was not assessed (Cheng et al., 2014). Given the lack of published studies addressing our concern, we compared three routinely used methods of preparing single cell suspensions of Mtb: low-speed spin, gentle sonication followed by low-speed spin, or filtration through a 5μm filter. We found that the method of bacterial preparation had a marked impact on intracellular bacterial viability, the global transcriptional pattern of infected cells, macrophage secretion of key innate immune mediators, and the ultrastructure of the bacterial cell envelope. Finally, when comparing an Mtb mutant that lacks PDIM to WT bacilli, we found that the method of preparation had a substantial impact on the inflammatory response to the mutant bacilli.

## Results

### Macrophage transcriptional responses to Mtb depend on the method of bacterial preparation

To investigate whether the method of dispersing bacterial cultures impacts host responses, we examined gene expression profiles of bone marrow-derived macrophages (BMDMs) that were uninfected or infected with WT Mtb (H37Rv strain) that were prepared either by passing through a 5μm filter (5μmF) or by brief sonication (so). The sonicated samples underwent three 10-second cycles in a water bath sonicator followed by a low-speed spin (sp), as described in **Methods**, and are designated so/sp. Bacilli prepared by the two methods were added to BMDMs at a multiplicity of infection (MOI) of 5, washed to remove extracellular bacteria after 4 h, and processed for RNA-seq 72 hours post-infection (hpi). We found that 536 genes were differentially expressed between uninfected cells and both of the infected samples (adjusted P value ≤0.01; fold change ≥|2|). Surprisingly, however, there were even more genes that were differentially expressed uniquely in macrophages infected by only one of the two bacterial preparations. 902 differentially expressed genes (DEGs) were unique when we compared uninfected with so/sp-infected macrophages, while 122 genes were uniquely differentially expressed in response to 5μmF bacteria (**Fig. 1A; Supplementary Figure 1A; Supplementary Table 1**). When we compared the DEGs in BMDM infected with so/sp-Mtb to those infected with the 5μmF-preparations, there were 732 DEGs (**Fig. 1A-B**). These included genes encoding important host defense molecules, including *Il6*, *Nos2*, and *Il1b*, which were markedly higher in so/sp-infected BMDMs compared to 5μmF-infected BMDMs (**Fig. 1B**).

**Figure 1.**
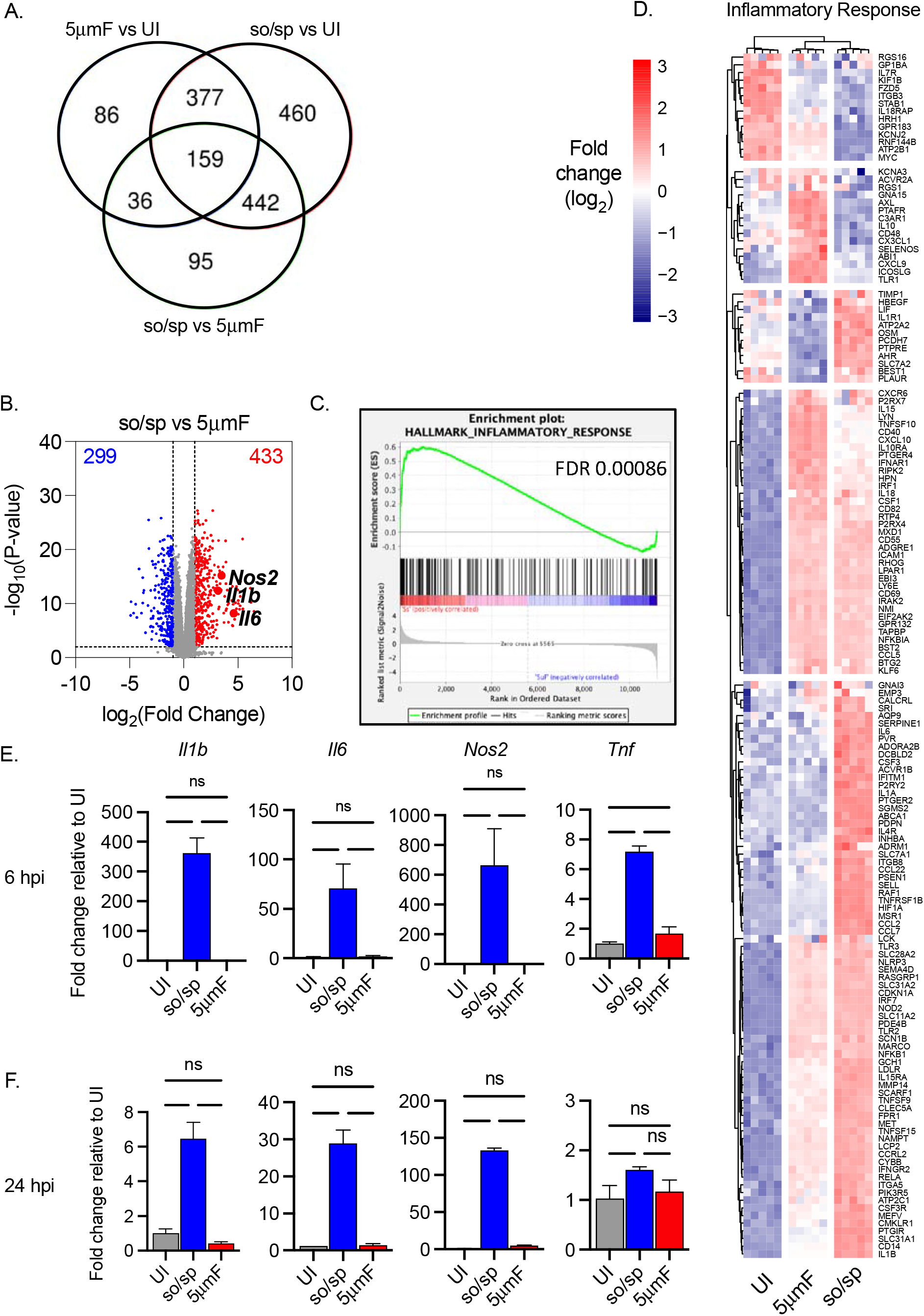
Cell preparation methods of Mtb impact macrophage responses. (**A-D**) BMDMs were uninfected or infected with Mtb prepared by sonication and spin (so/sp) or filtration (5μmF) at an MOI of 5 and analyzed 72 hpi by RNA-seq (n=5 per condition). **(A)** Venn diagram illustrates the number of DEGs between samples. **(B)** Volcano plot shows genes differentially expressed in BMDMs infected with so/sp versus 5μmF Mtb. DEGs exhibiting an adjusted P-value of ≤0.01 and a linear fold change ≥2.00 (red) or ≤2.00 (blue) are indicated. **(C-D)** GSEA identified hallmark gene sets that were significantly enriched in so/sp-versus 5μmF-infected BMDMs (P ≤0.01; FDR ≤0.01). Representative enrichment plot **(C)** and corresponding heat map **(D)** for the gene set “inflammatory response.” Expression values in heatmap were generated using log2 normalized CPM for each gene. **(E and F)** qPCR was performed on uninfected BMDMs or BMDMs infected with so/sp- or 5μmF-prepared Mtb 6 **(E)** and 24 **(F)** hpi using an MOI of 10. Data are shown as fold change in gene expression relative to uninfected BMDMs. Data shown are mean +/− SD from one representative experiment with three biological replicates per group and two technical replicates per sample. qPCR experiments were performed at least 3 independent times. Statistical significance was determined with one-way ANOVA using Tukey’s multiple comparisons test. **(A-F)** Error bars indicate mean +/− SD. ns not significant; *P<0.05; ***< 0.001; ****< 0.0001.

In order to further analyze the transcriptional differences, we used Gene Set Enrichment Analysis (GSEA) to query our expression data against hallmark gene sets from the Molecular Signatures Database (Liberzon et al., 2015). We found that 10 hallmark gene sets were significantly enriched in so/sp preparations relative to 5μmF-infected BMDMs (P ≤0.01; FDR ≤0.01) (**Fig. 1C-D, Supplemental Fig. 1B**). The sonicated bacilli elicited a significant enrichment of gene sets that included TNFA signaling via NFKB, inflammatory response, MTORC1 signaling, glycolysis, xenobiotic metabolism, and IL6 JAK STAT3 signaling (**Fig. 1C, Supplemental Fig. 1B**). In contrast, 1 hallmark gene set was significantly enriched in 5μmF-relative to so/sp-infected BMDMs (E2F targets) (**Supplemental Fig. 1C**). Visualization of transcriptional data from the hallmark gene set “inflammatory response” showed a distinct gene expression pattern in response to so/sp versus 5μmF bacteria (**Fig. 1D**). Overall, the macrophages infected with the so/sp bacilli displayed a more robust pro-inflammatory phenotype, whereas the 5μmF-infected macrophages were enriched in pro-replication pathways. In addition to the 72 hpi timepoint used for RNA-seq, we found that infection with so/sp Mtb elicited significantly higher levels of expression of *Il1b*, *Nos2*, *Il6* and *Tnf* at 6 and 24 hpi by qPCR compared with 5μmF preparations, with the greatest difference seen early in infection (**Fig. 1E-F**). Strikingly, while the so/sp bacilli markedly upregulated inflammatory gene expression, there was minimal difference between 5μmF-infected and uninfected macrophage 6 hpi and 24 hpi. In these experiments, we used 5μm filters with hydrophilic polyethersulfone (PES) membranes, which are composed of aryl-SO_2_-aryl subunits. We considered the possibility that the chemical backbone and hydrophilic nature of the filter might be altering Mtb, but we had similar findings when we used hydrophobic polytetrafluoroethylene (PTFE) filters (**Supplemental Fig. 2**). In conclusion, the transcriptional response of BMDMs to Mtb infection was markedly different depending on the method of bacterial preparation.

### Sonication increases the inflammatory impact of Mtb

Given the dramatic difference between so/sp and 5μmF bacilli, it was important to assess which one more accurately reflects unperturbed Mtb. However, if we were to use Mtb directly from a liquid culture, it would not be possible to establish that we are using similar numbers of bacilli compared to the other preparations given the propensity to clump. Therefore, we used a low-speed spin preparation. This was the same procedure applied to the so/sp sample, but the sonication step was omitted. Specifically, liquid cultures were centrifuged at 206 × g for 10 minutes, after which the supernatant was removed and centrifuged at 132 × g for 8 minutes, and the final supernatant was used to infect BMDMs. We found that macrophages infected with the spin (sp) sample had an intermediate phenotype between so/sp and 5μmF samples (**Fig. 2A**), eliciting significantly less *Il1b*, *Il6*, *Nos2*, and *Tnf* expression than the so/sp samples. To determine whether the impact of preparation method was specific to H37Rv, we tested two additional Mtb strains: HN878, a W-Beijing lineage strain that was isolated in a TB outbreak in Houston in the 1990s, and Erdman, a strain that is commonly used is laboratory studies. We found that the method of preparation had a similar impact on macrophage inflammatory responses for these strains as for H37Rv (**Supplemental Fig. 3**). A variety of Mtb PAMPs have been shown to activate TLR2 (Hinman et al., 2021). In order to establish whether the so/sp samples were activating TLR2-dependent pathways, we infected BMDMs from *Tlr2^−/−^* mice. We found that expression of *Il1b*, *Il6*, *Nos2*, and *Tnf* were significantly reduced in TLR2 KO BMDMs in response to so/sp Mtb relative to WT BMDMs (**Fig. 2B**). This was also true for the induction observed in response to spin preparations.

**Figure 2.**
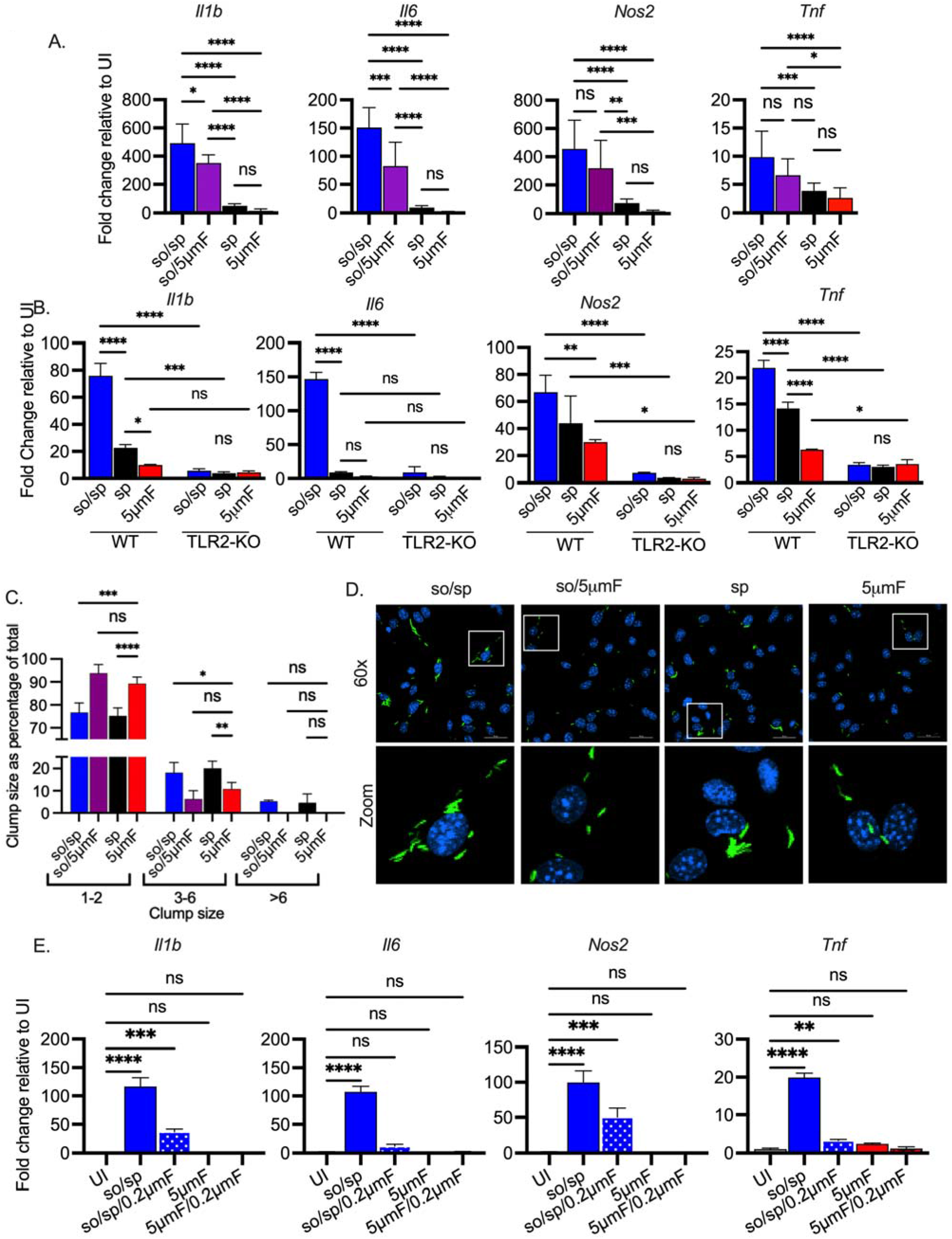
Sonicated bacteria induce high TLR2-dependent inflammatory responses. **(A)** BMDMs were uninfected or infected with different preparations of Mtb as indicated at an MOI of 10 and analyzed by qPCR 6 hpi. Data are presented as fold changes in gene expression relative to uninfected BMDMs. Data are combined from 2 to 3 experiments, each with three biological replicates per group and two technical replicates per sample. Statistical significance was determined with one-way ANOVA using Tukey’s multiple comparisons test. **(B)** WT or *Tlr2*^−/−^ BMDMs were uninfected or infected with different preparations of Mtb as indicated for 6 h at an MOI of 10 and analyzed by qPCR. Data are presented as fold change in gene expression relative to uninfected BMDMs of the same mouse genotype. Data are representative of 3 experiments, each with three biological replicates per group and two technical replicates per sample. Statistical significance was determined with two-way ANOVA using Tukey’s multiple comparisons test. **(C)** BMDMs were infected with GFP-expressing bacteria (MOI 5) and visualized using immunofluorescence at 4 hpi. Bacteria were quantified and classified as single/doublets (1-2), small (3-6), or large (>6) clumps and quantified for each preparation. At least 100 bacterial occurrences of each preparation method were analyzed. Two-way ANOVA with Dunnett’s multiple comparisons test was used to assess statistical significance within each batch relative to the given 5μmF quantitation. **(D)** Representative fluorescence microscopy images of BMDMs (nuclei stained with DAPI) infected with GFP-expressing Mtb used in (C). Images are maximum-intensity projections. Boxed areas in the merged image are shown in higher magnification in the bottom panel. **(E)** BMDMs were untreated, infected with the indicated bacterial preparations at MOI 10, or treated with the sterile filtrate from different preparations for 6h and analyzed by qPCR. Data are presented as fold changes in gene expression relative to untreated BMDMs. Data are representative of 3 experiments, each with three biological replicates per group and two technical replicates per sample. Statistical significance was determined for each group relative to untreated BMDMs with one-way ANOVA using Dunnett’s multiple comparisons test. **(A-B, D)** Error bars indicate mean +/− SD. ns not significant; *P<0.05; **<0.01; ***< 0.001; ****< 0.0001.

We wondered if the 5μmF bacilli contained a factor that inhibited macrophage gene expression or if they were just less proinflammatory. To address this, we mixed so/sp and 5μmF bacilli together in equal proportions and assayed gene expression by qPCR. We found that the mixed samples were still inflammatory, arguing against a potent inhibitory factor coming from the filtered preparation (**Supplemental Fig. 4A**). In addition, if we prepared bacteria by first sonicating and then using a 5μmF (so/5μmF), the expression changes resembled so/sp infection, with marked upregulation of *Il1b*, *Il6*, *Nos2*, and *Tnf* (**Fig. 2A**). We considered the possibility that the different inflammatory responses might be a result of different degrees of aggregation of the bacilli in each preparation. To visualize the bacteria, we infected BMDMs with GFP-expressing Mtb prepared by the various methods and examined them by fluorescence microscopy (**Fig. 2C-D**). We quantified whether the visualized bacteria were single/doublets (1-2), small (3-6), or large (>6) clumps. For all of the preparations, more than 75% of the bacterial occurrences were single/doublets. The 5μmF and so/5μmF preparations had slightly more single/doublets and slightly fewer clumps than the other samples (**Fig. 2C-D**). Since the clumpiness of so/sp and sp samples were similar, aggregation status did not explain the hyper-inflammatory nature of the so/sp samples. In addition, when the sonicated sample was filtered (so/5μmF), it had few clumps, and the bacilli still induced high levels of *Il1b*, *Il6*, *Nos2*, and *Tnf* (**Fig. 2A-D**). To investigate whether the response to sonicated bacteria was due to soluble factors released from the bacilli, we passed the so/sp sample through a 0.2 μm filter to remove bacteria. We treated macrophages with equal volumes of the sterile filtrate or the unfiltered so/sp sample and analyzed subsequent gene expression. In support of extra-bacterial components contributing to the inflammatory gene expression, the expression of *Il1b, Nos2, Il6, and Tnf* were all significantly increased in response to the sterile filtrate prepared from the so/sp bacteria compared to uninfected BMDMs (**Fig. 2E**). In contrast, there was no difference in expression of these genes in the sterile filtrate of sp or 5μmF bacteria relative to uninfected BMDMs (**Fig. 2E**, **Supplemental Fig. 4B**). To conclude, compared to bacteria prepared by a low-speed spin or 5μmF, bacilli that were sonicated induced substantially higher TLR2-dependent transcriptional responses in macrophages, independent of their aggregation status and due in part to soluble mediators.

To determine whether the changes in gene expression resulted in altered cytokine secretion, we used the FluoroDOT assay to evaluate secretion of TNF-α. This approach uses plasmon-enhanced fluorescent nanoparticles called plasmonic fluors to visualize protein secretion by microscopy (**Supplemental Fig. 5**). This allowed us to examine secretion of TNF-α at an early time point after infection and with single cell resolution (Seth et al., 2022). Similar to the transcriptional data, the sonicated preparations elicited the most TNF-α secretion followed by the sp and 5μmF preparations (**Fig. 3A-B**). We confirmed these findings by measuring TNF-α by enzyme linked immunosorbent assay (ELISA; **Fig. 3C**). We also evaluated IL-1β secretion using ELISA and found that the so/sp preparation elicited increased secretion of IL-1β **(Fig. 3C**). Interestingly, we found that when infected by the sonicated samples, most of the macrophages, both infected as well as uninfected bystanders in the same well, secreted TNF-α. In contrast, infection with the sp or 5μmF Mtb resulted in only infected cells secreting TNF-α (**Fig. 3A**). In addition, so/sp samples that had been sterilized by passage through a 0.2 μm filter elicited significantly more TNF-α secretion than sterilized 5μmF samples (**Fig. 3D-E**). This is consistent with the observation that extra-bacterial components in the sonicated preparation contribute to inflammatory gene expression.

**Figure 3.**
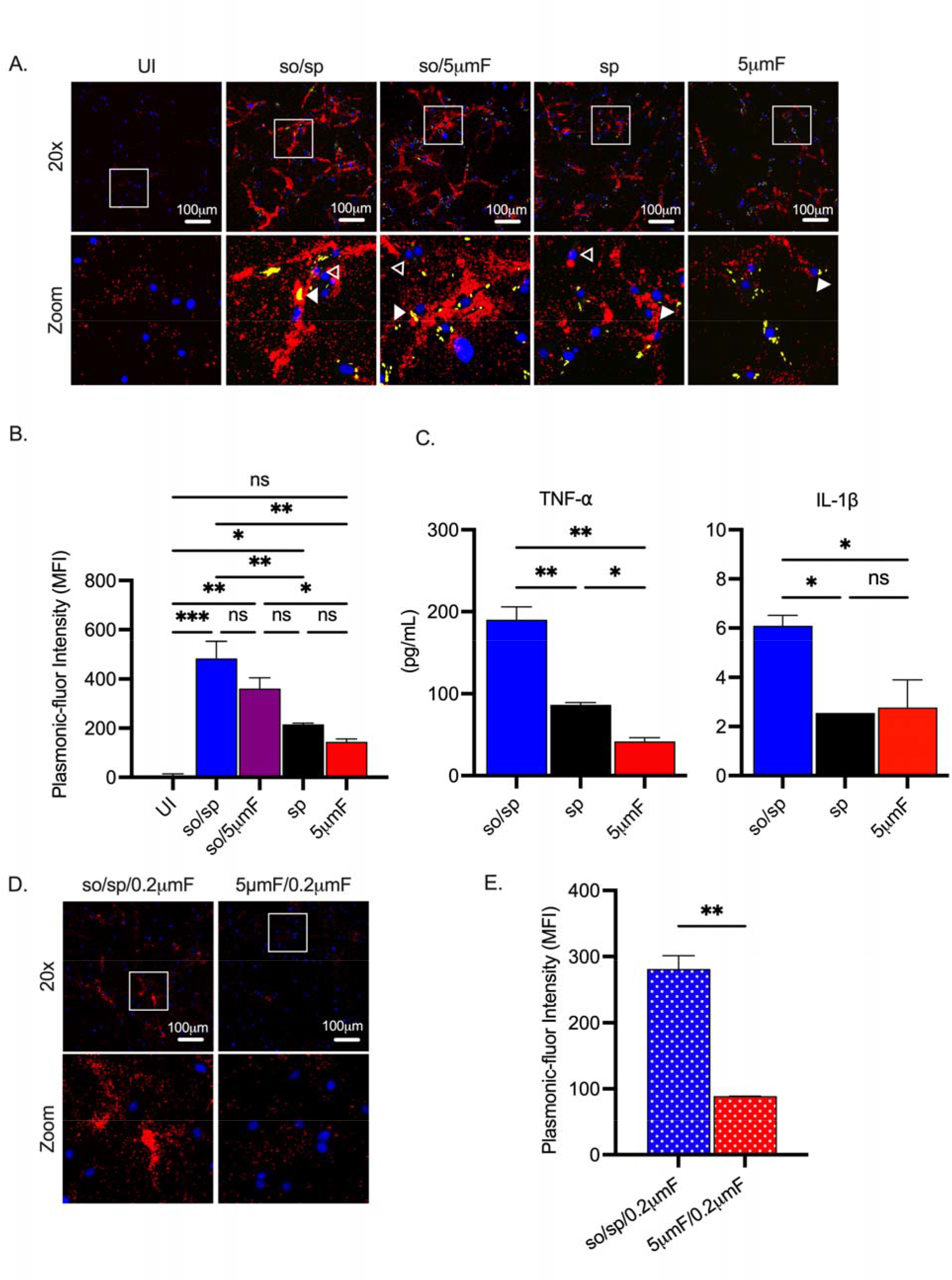
Sonicated bacteria elicit elevated TNF-α and IL-1β secretion. **(A)** Using the FluoroDOT assay, BMDMs were grown on a glass bottom plate that was coated with TNF-α capture antibody, infected at an MOI of 10 with H37Rv-GFP prepared by the indicated method, and examined by epifluorescence microscopy (20X) 6 hpi. Images show Plasmonic-fluor 650 (red), Mtb (GFP), and DAPI (blue). Boxed areas in the image are enlarged in the bottom images. Secretion from infected BMDMs or uninfected bystander cells are highlighted by open or closed white arrowheads, respectively. **(B)** Data show the quantification of the mean fluorescence intensity (MFI) of the plasmonic-fluor in the entire well from each different condition shown in **(A)**, with statistical significance determined with one-way ANOVA using Tukey’s multiple comparisons test. **(C)** IL1-β and TNF-α were measured 24 hpi in the culture supernatant of uninfected or Mtb-infected BMDMs (MOI 10) by ELISA. Data shown are mean +/− SD from one representative experiment with three biological replicates per group and two technical replicates per sample. Significance was determined using one-way ANOVA with Tukeys’ multiple comparisons test. **(D)** Using the FluoroDOT assay, BMDMs grown on a glass bottom plate that was coated with TNF-α capture antibody were exposed to the sterile filtrate of bacterial single cell suspension prepared by either so/sp or 5μmF and examined by epifluorescence microscopy (20X) 6 hpi. Images show Plasmonic-fluor 650 (red) and DAPI (blue). Boxed areas in the image are enlarged in the bottom images. **(E)** Data show the quantification of the mean fluorescence intensity (MFI) of the plasmonic-fluor in the entire well from each condition shown in D, with statistical significance determined using an unpaired T test. **(A-E)** Error bars indicate mean +/− SD. ns not significant; *P<0.05; **<0.01; ***<0.001.

### Filtered Mtb are attenuated in BMDMs

Given that the different preparations generated pronounced differences in macrophage gene expression and cytokine secretion, we hypothesized that they would also exhibit differences in intracellular viability. We infected BMDMs with Mtb prepared by the different methods. To ensure that a similar MOI was used for each bacterial preparation, we plated the input used for the infection. We had to use 1.5-times more filtered (so/5μmF and 5μmF) bacilli based on OD_600_ to achieve the same number of viable bacteria. After infection, the intracellular bacilli were enumerated at 4 hpi and 3 and 5 days post-infection (dpi). Mtb that were prepared by so/sp or sp grew in macrophages significantly better than those that were filtered (so/5μmF and 5μmF; **Fig. 4A**). Increasing the MOI of the filtered bacteria to 20 or 40 did not overcome the intracellular growth defect (**Fig. 4B**). The differences in intracellular growth were not explained by differences in macrophage viability (**Fig. 4C**). We verified that the filtered Mtb were still viable, as they grew indistinguishably from other preparations when they were inoculated in liquid culture (**Fig. 4D**). Similar to our findings with H37Rv, filtered HN878 and Erdman were also attenuated in BMDMs compared to those prepared by so/sp or sp (**Fig. 4E-F**). Thus, filtered Mtb appeared to be both less inflammatory and impaired in their ability to counter the antimicrobial properties of BMDMs.

**Figure 4.**
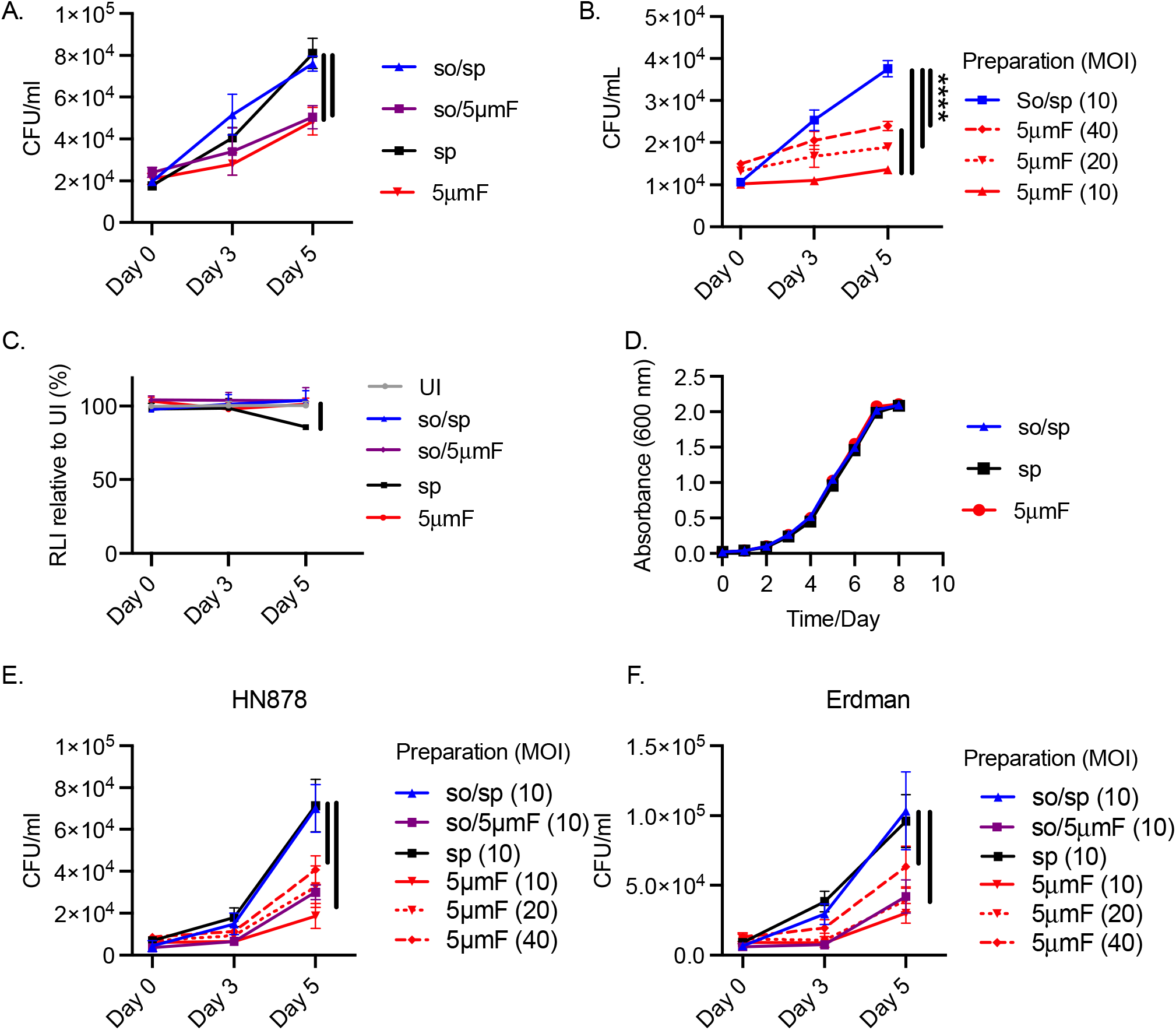
Filtered Mtb are attenuated in BMDMs. **(A)** BMDMs were infected with different preparations of Mtb (H37Rv) at an MOI of 10 and intracellular bacteria were enumerated by colony forming units (CFU) 4 hpi, 3 dpi, or 5 dpi. **(B)** BMDMs were infected with different preparations of Mtb (H37Rv) at an MOI of 10-40 and intracellular bacteria were enumerated by CFU 4 hpi, 3 dpi, or 5 dpi. **(C)** BMDMs were infected with different preparations of Mtb (H37Rv) at MOI of 10, and BMDM viability was measured using the CellTiter-Glo assay at 4 hpi, 3 dpi, and 5 dpi. Statistical significance between preparations was determined with two-way ANOVA using Dunnett’s multiple comparisons test with selected significance values presented for 5 dpi relative to UI BMDMs. **(D)** Growth curve of different bacterial H37Rv preparation in liquid media (7H9 media supplemented with 10% Middlebrook OADC, 0.05% Tyloxapol, and 0.2% glycerol). **(E-F)** BMDMs were infected with different preparations of HN878 (**E**) or Erdman (**F**) strains at an MOI of 10-40 and intracellular bacteria were enumerated by CFU 4 hpi, 3 dpi, or 5 dpi. **(A-C, E-F)** For all CFU and macrophage viability studies, six biological replicates were used per group. Statistical significance was determined for each preparation at 5 dpi by comparing to CFU from BMDMs infected with spin-prepared Mtb using two-way ANOVA with Dunnett’s multiple comparisons test. Selected significance values are presented at 5 dpi. **(A-F)** Error bars indicate mean +/− SD. ns not significant; ****<0.0001.

### Sonication and filtering affect the bacterial cell wall

To determine if there were structural differences between the sonicated, spun, and filtered Mtb, we used transmission electron microscopy (TEM). We first generated ultrathin cross-sections of bacteria to visualize the ultrastructure of the cell envelope (**Fig. 5A-C**). In bacteria prepared with low-speed spin, we could distinguish the structural layers of the cell envelope that have been previously described: the innermost phospholipid bilayer, followed by electron-dense peptidoglycan and arabinogalactan layers, a translucent mycobacterial outer membrane, and an outermost carbohydrate-rich capsular layer (**Fig. 5B)**. In bacteria prepared with low-speed spin and/or sonication, each of these distinct layers were apparent (**Fig. 5A-B**). In the 5μmF-prepared bacteria, the phospholipid bilayer was seen, surrounded by an electron dense layer, but there appeared to be loss of the capsular layer and potentially the mycomembrane as well (**Fig. 5C**). While TEM of ultrathin cross-sections provided excellent resolution of the cell wall, it was also subject to artifact introduced by drying and fracturing of the bacteria required in this technique. This made it difficult to know how representative the well-preserved bacilli were in terms of the total population. Therefore, we also visualized bacilli by adsorption to a copper grid followed by 1% uranyl acetate staining, a simple technique which minimized artifact (**Fig. 5D-F**). Uranyl acetate is a common negative stain used for TEM that can bind to capsular polysaccharides (Stukalov et al., 2008), and it created an electron dense halo around the bacteria. We quantified the width of the electron dense halo on individual bacilli and found that it was significantly thinner on bacteria prepared with the 5μmF compared to so/sp and sp (**Fig. 5G**). This suggests a different chemical composition of the outermost layer of the filtered bacteria and was consistent with the differences noted in the TEM. In addition, the samples from 5μmF-treated bacteria had substantial extracellular debris, which may be damaged fragments from the outer layers of the envelope. Finally, more dead bacteria were noted in the 5μmF sample as evidenced by penetration of the dark staining uranyl acetate into the cells **(Fig. 5F)**, which may explain why we had to use 1.5-times more 5μmF bacilli (based upon optical density) to achieve the same number of viable bacteria. Using this technique, we also observed that the so/sp bacteria, but not sp or 5μmF bacteria, had prominent round protuberances that were approximately 0.2 to 1 μM in diameter present on the outer surface of the bacteria or, less frequently, in the culture media **(Fig. 5D)**. To conclude, both sonicated and filtered preparations had evidence of distinct types of damage to the envelope on TEM that were not apparent in the samples which had been prepared by centrifugation alone.

**Figure 5.**
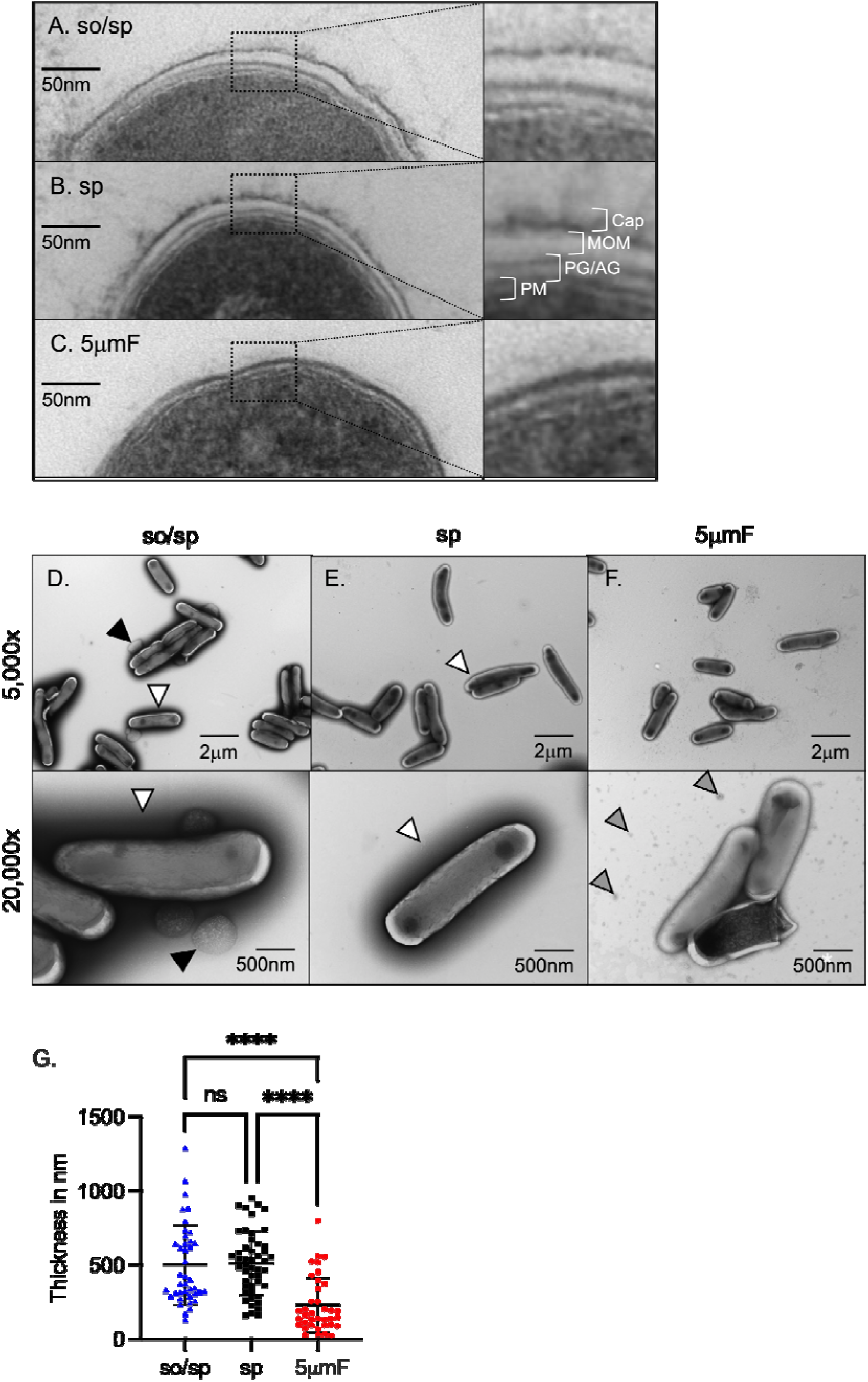
Sonication and filtering affect the bacterial cell wall. **(A-C)** TEM of ultrathin cross-sections of Mtb at 50,000x magnification (left) beside enlarged cross-section of the envelope (right). The plasma membrane (PM), peptidoglycan/arabinogalactan layer (PG/AM), mycobacterial outer membrane (MOM), and capsular layer (Cap) are indicated. **(D-F)** Mtb were absorbed on freshly glow discharged formvar/carbon-coated copper grids followed by negative staining with 1% aqueous uranyl acetate. Representative images are 5,000x (above) and 20,000x (below). So/sp-prepared Mtb had round protuberances that were on or near their envelopes indicated by black arrows. Electron-dense outer halos seen surrounding so/sp- and sp-prepared bacteria are indicated with white arrows. Debris seen in the extracellular space of 5μmF-prepared Mtb is indicated with gray arrows. **(G)** Capsule thickness was measured in nanometers using TEM images from bacteria stained with 1% uranyl acetate. Thickness measurements were compared between preparations with one-way ANOVA using Tukey’s multiple comparisons test. Error bars indicate mean +/− SD. ns not significant; ****<0.0001.

### The interpretation of the role of PDIM in inflammatory responses depends upon preparation method

PDIM is a multifunctional virulence lipid that is present in the envelope of members of the Mtb complex as well as closely related *Mycobacterium marinum*. Along with the ESX-1 type VII secretion system, PDIM facilitates phagosomal escape of Mtb, a crucial event that allows the bacteria to gain access to the cytosol, subvert cell death pathways, and promote extracellular spread (Augenstreich et al., 2017; Barczak et al., 2017; Cox et al., 1999; Lerner et al., 2018; Osman et al., 2020; Quigley et al., 2017). In addition, PDIM contributes to the low permeability of the mycobacterial envelope, alters the host’s initial innate immune response, and may physically shield mycobacterial PAMPs or interfere with their activation of PRRs (Astarie-Dequeker et al., 2009; Camacho et al., 2001; Cambier et al., 2014; Murry et al., 2009; Rousseau et al., 2004; Siméone et al., 2007). To determine whether PDIM dampens inflammatory signaling, we used a strain with a deletion in *ppsD*, which results the in the absence of PDIM (Barczak et al., 2017). When we examined macrophage gene expression after infection with Δ*ppsD* by qPCR, we found that expression of *Il1b*, *Il6*, *Nos2*, and *Tnf* was significantly increased compared to infection with WT Mtb, consistent with the idea that PDIM reduces inflammatory signaling (**Fig. 6A**). However, this was only significant and reproducible in the sonicated sample; there was little difference between Δ*ppsD* and WT Mtb if they were prepared by sp or 5μmF. We had similar findings when we used the FluoroDOT assay to examine TNF-α secretion (**Fig. 6B-C**). The Δ*ppsD* mutant reproducibly elicited more TNF-α secretion than WT Mtb, but only if the sample was sonicated. Interestingly, when we examined the Δ*ppsD* mutant by TEM, we found that the Δ*ppsD* mutant lacked the dark halo that was seen in so/sp and sp samples of WT Mtb; the halo was restored by complementation, suggesting that lack of PDIM altered the interaction of uranyl acetate with the mycobacterial surface (**Fig. 6D-I**). This difference, however, is unlikely to account for the hyperinflammatory signaling, as it was seen in all Δ*ppsD* samples, and only the sonicated samples were hyperinflammatory. As we had seen with WT Mtb, there were round protrusions and vesicles in sonicated sample of both Δ*ppsD* and the complemented strain **(Fig. 6D, G)**. There was no obvious visual difference between the Δ*ppsD* mutant and complemented strain to explain why the so/sp Δ*ppsD* mutant was more hyperinflammatory than so/sp WT. To conclude, the hyperinflammatory phenotype associated with the Δ*ppsD* mutant depended upon the method of bacterial preparation.

**Figure 6.**
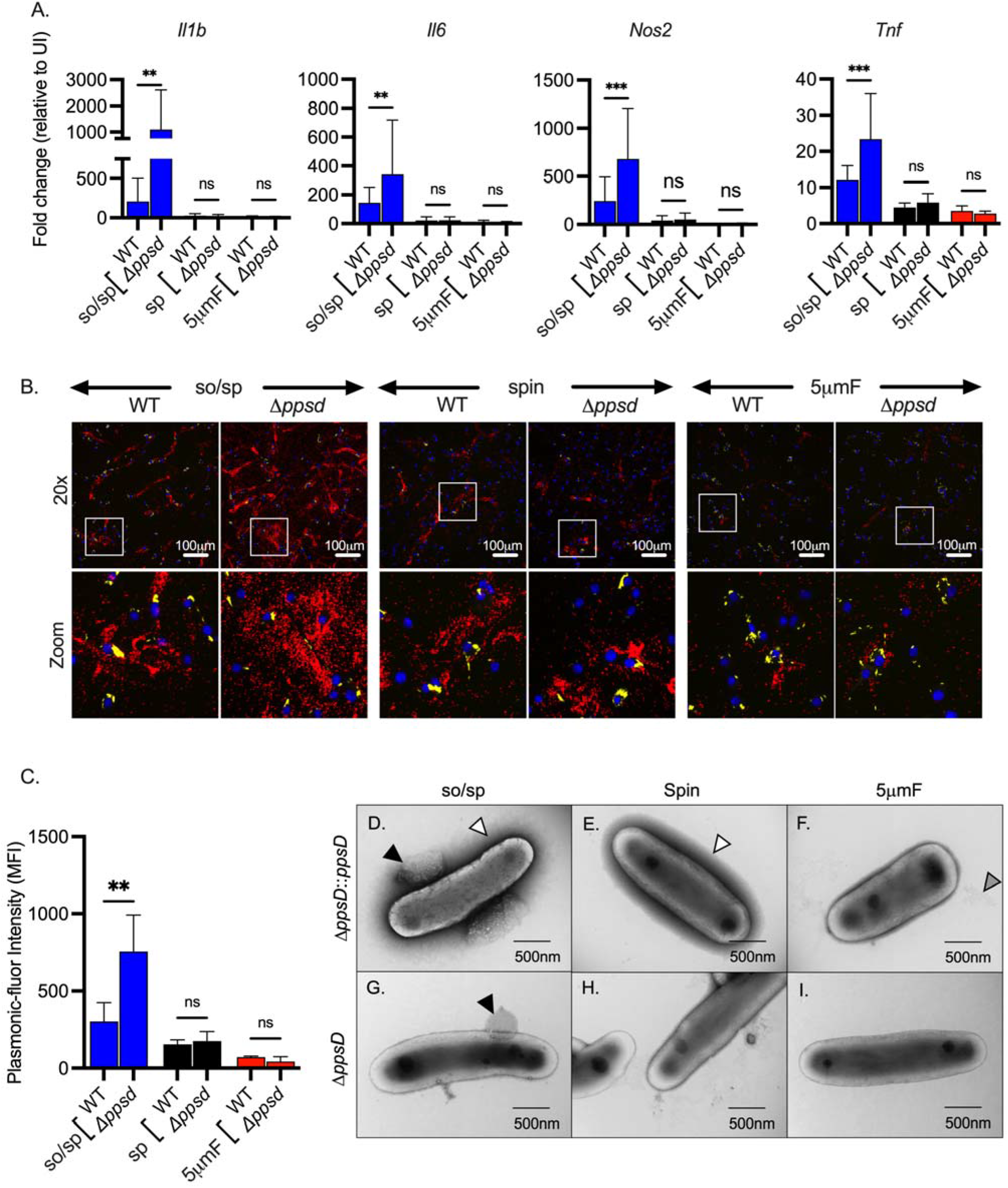
The role of PDIM in inflammatory responses depends upon preparation method. **(A)** BMDMs were uninfected or infected with indicated strains of Mtb at an MOI of 10, and gene expression was analyzed by qPCR at 6 hpi. Data are presented as fold change in gene expression relative to uninfected BMDMs of the same mouse genotype. Statistical significance was determined with two-way ANOVA using Tukey’s multiple comparisons test. Data are combined from 2 to 3 experiments, each with three biological replicates per group and two technical replicates per sample. **(B)** Using the FluoroDOT assay, BMDMs were grown on a glass bottom plate that was coated with TNF-α capture antibody, infected at an MOI of 10 with H37Rv-GFP or Δ*ppsD*-GFP prepared by the indicated method, and examined by epifluorescence microscopy (20X) 6 hpi. Images show Plasmonic-fluor 650 (red), Mtb (GFP), and DAPI (blue). Boxed areas in the image are enlarged in the bottom images. **(C)** Data show the quantification of the mean fluorescence intensity (MFI) of the plasmonic-fluor in the entire well from each conditions shown in B with statistical significance determined with two-way ANOVA using Tukey’s multiple comparisons test. **(D)** Bacteria were imaged by allowing indicated Mtb strains to absorb on freshly glow discharged formvar/carbon-coated copper grids followed by negative staining with 1% aqueous uranyl acetate. Round protuberances seen on or near the envelopes of so/sp-prepared H37Rv Mtb are indicated by black arrows, the electron-dense outer halos seen surrounding so/sp- and sp-prepared H37Rv Mtb are indicated with white arrows, and the debris seen in 5μmF-prepared H37Rv Mtb are indicated with gray arrows. **(A, C)** Error bars indicate mean +/− SD. ns not significant; **P<0.01; ***<0.001.

## Discussion

More than fifty years ago, D’Arcy Hart demonstrated that *M. tuberculosis* avoids lysosomal delivery within macrophages (Armstrong & Hart, 1971). Ever since his landmark study, in an effort to understand fundamental mechanisms of TB pathogenesis, investigators have studied the interaction of Mtb with macrophages. They have also employed a variety of methods to disperse the bacilli to enable subsequent analysis. Unexpectedly, we found that two commonly used single cell preparation methods significantly impacted Mtb-host interactions: sonicated bacilli were hyperinflammatory, and 5μm-filtered Mtb were attenuated in macrophages. These effects were seen for H37Rv, HN878, and Erdman strains. In addition, we found that the method of preparation changes the impact of PDIM on the early macrophage transcriptional responses to Mtb. Consistent with the data of Hinman *et al*, who used a low-speed spin method to remove clumps (Hinman et al., 2021), we found little impact of PDIM on the early TLR2-dependent response to centrifuged bacteria. However, if the bacteria were briefly sonicated, then strains without PDIM elicited increased pro-inflammatory gene expression compared to control strains. This suggests that the mutant without PDIM is either more sensitive to sonication-induced damage or that it is more inflammatory once that damage occurs. It is important to point out that we only examined the early TLR2-dependent response. PDIM influences a variety of processes, including later TLR2-driven responses, phagosomal escape, and intracellular survival (Augenstreich et al., 2017; Barczak et al., 2017; Cambier et al., 2020; Hinman et al., 2021; Lerner et al., 2018; Osman et al., 2020; Quigley et al., 2017). It is possible that these other PDIM-dependent processes are not impacted by the preparation method. Nonetheless, our studies demonstrate that the preparation method needs to be considered in host-pathogen interaction studies, as it can change the interpretation of bacterial mutants and has a dramatic effect on TLR2-dependent responses and intracellular bacterial survival in bone marrow-derived macrophages.

Given the extensive use of macrophages in Mtb pathogenesis studies, there are surprisingly few studies investigating the impact of dispersal methods. A 2004 study demonstrated that prolonged sonication (5 minutes) reduces Mtb viability, while bacteria that had undergone gentle sonication (30 sec × 3) exhibited enhanced binding to macrophages and altered surface charge relative to syringed bacteria (Stokes et al., 2004). In that study, the sonicated bacteria had an altered cell envelope, which appeared uneven and bulging. Even though we sonicated for a shorter time (10 sec × 3), we also saw evidence of similar cell envelop disruption by TEM. In addition, we found that sonicated bacteria elicited orders-of-magnitude higher levels of TLR2-dependent transcriptional responses, leading to enhanced IL-1β and TNF-α secretion. This was mediated in part by material that was no longer cell associated, as even sterile-filtered samples activated macrophages. In addition, uninfected bystander cells that had been treated with so/sp preparations secreted TNF-α in the FluoroDOT assay. Our TEM findings suggest that sonication results in cell envelope damage and generation of small structures that resemble extracellular vesicles (EVs) that have been described in Mtb, although the vesicles that we saw are generally larger than the majority of EVs (Prados-Rosales et al., 2011). Mtb EVs are formed by an active process and contain immunomodulatory molecules including lipoarabinomannan and other TLR2 agonists (Athman et al., 2015; Palacios et al., 2021; Prados-Rosales et al., 2011). Whether the structures formed by sonication have similar content to EVs found in growing cultures will require further studies.

How might sonication and filtration lead to the distinct macrophage responses that we observed? Electron microscopy revealed that sonication and filtration cause different types of alteration to the cell envelope (**Supplemental Fig. 6A**). The cell envelope is an elaborately layered structure that contains a variety of lipid and protein PAMPs and virulence factors. Our data are consistent with a model in which PAMPs and virulence factors are differentially impacted by sonication and filtration (**Supplemental Fig. 6B**). In this model, sonication disrupts the cell envelop in a manner that makes PAMPs more highly exposed or accessible, while leaving the activity of virulence factors intact. In contrast, filtration (5μmF) disrupts the cell envelop in such a way that both virulence factors and PAMPs are inactivated and/or dispersed from the bacilli, rendering the bacteria both attenuated and less inflammatory. Bacteria that are subject only to low-speed spin are neither hyper-inflammatory nor attenuated, since both PAMPs and virulence factors are less perturbed. In the case in which the bacteria are sonicated and filtered, they are both hyper-inflammatory and attenuated, which can be explained by enhanced exposure of PAMPs through sonication and inactivation of virulence factors by filtration.

This is one model that would explain our findings, but other scenarios could be envisioned. Mtb cultures are highly heterogenous, so we considered the possibility that a small minority of the so/sp bacilli were contributing to the heightened inflammatory response. However, the Fluoro-dot data, which allows us to visualize cytokine secretion at a single cell level, argue against this possibility. Similarly, if there were a small population of bacilli in the filtered sample that were inhibiting the inflammatory response, then, we would have expected the mixed samples to behave like the filtered sample, but rather we saw an intermediate phenotype. In terms of intracellular growth, there is undoubtedly heterogeneity within the population, with better intracellular growth on a population level in the so/sp and sp samples relative to filtration, with all samples having a mix of growth, stasis, and killing. Two aspects of heterogeneity that we assessed are clump size and capsule thickness. On a population level, we found a significant reduction in the thickness of the capsule of the 5μmF bacteria compared to both so/sp and sp bacteria. However, there was a wide distribution in cell envelope thickness, which may contribute to heterogeneity in macrophage responses to bacilli on a cell-to-cell level. Finally, we considered that differences in the degree of aggregation in the different bacterial preparations may account for differences in inflammatory potential or intracellular survival (Rodel et al., 2021), but heterogeneity in this aspect of the bacterial population was unlikely to explain the differences in outcomes. There may be other aspects of underlying heterogeneity in our samples that contribute to their distinct behavior or confound bulk measurements.

While our study was limited to three common single cell preparation methods, we expect that other techniques would also impact the mycobacterial envelope and host interactions. We queried PubMed for papers published in 2021 on Mtb and macrophages to determine which methods were commonly used (**Supplemental Fig. 7**). Of the 119 papers, only 39.5% reported how they generated single cell suspensions. Of those that did report their methodology, 42.5% used more than one method. The most commonly reported methods were syringing, followed by sonication and low-speed centrifugation. Less often, filtering, vortexing with glass beads, or allowing gravity to sediment the larger clumps were used. Dispersing clumps with glass beads would likely disrupt the envelope, as studies have used this technique to selectively remove and isolate the capsular layer to analyze its components (Lemassu & Daffe, 1994; Lemassu et al., 1996; Ortalo-Magne et al., 1995). Others have reported that syringing through a 25-gauge needle produced no apparent disruption to the envelope on TEM (Stokes et al., 2004), but these samples were not evaluated further in terms of macrophage responses. We did not evaluate syringing, because it is not an approved method in our biosafety level 3 facility due to the risk of aerosolization and needle stick injuries. The physical forces used to disrupt clumps by this method might also result in envelope alterations, and investigators should consider this in their studies. Overall, we consider centrifuged samples as the least disrupted, but even centrifugation might disrupt the capsule, as do detergents that are commonly used in liquid cultures (and were used for all of our studies). Detergents are known to cause release of capsular components into the culture filtrate (Kalscheuer et al., 2019; Sani et al., 2010), although the impact on host interactions is relatively unexplored. The impact of detergent treatment on cytokine responses and vaccine responses have been evaluated, but after detergent treatment, single cell suspensions were generated by filtering, sonicating, or syringing (Prados-Rosales et al., 2016; Sani et al., 2010), complicating the interpretation.

Even if investigators had a non-disruptive way to isolate single cells, the behavior of single cells may not be the same as large aggregates that make up a substantial fraction of the unperturbed bacterial population. The aggregation state of Mtb has long been reported to be important to pathogenesis. For example, the observation that Mtb forms serpentine cords *in vivo* dates back to the earliest descriptions of the bacteria. Aggregated Mtb are found at the periphery of human necrotic granulomas, in alveolar macrophages of Mtb-infected patients, and are exhaled by infected individuals (Dinkele et al., 2021; Rodel et al., 2021; Ufimtseva et al., 2018). The literature describing the impact of aggregation on host interactions is difficult to interpret in light of our findings, as many of these studies used agitation with glass beads, filtration, sonication, or some combination of these procedures to generate the dispersed samples (Kolloli et al., 2021; Mahamed et al., 2017; Rodel et al., 2021). Thus, bacterial aggregation is likely an important virulence property of Mtb, which investigators overlook in the effort to generate single cell suspensions; at the same time, in generating single cell suspensions, investigators introduce the potential for experimental artifact.

It is possible that technical differences in how other laboratories sonicate or filter bacteria could result in findings that are different from ours. We used log phase cultures of H37Rv, HN878, and Erdman strains that had been grown with gentle agitation, a fatty acid source (oleic acid), and 0.05% Tyloxapol to infect mouse macrophages, but other investigators use different strains, frozen stocks, omit oleic acid, use different detergents, or infect human macrophages, all of which could lead to differences from our findings. An important conclusion of our findings is that investigators should fully report the methods that they use to grow and process mycobacteria and consider the impact of the methodology on their findings.

While sonication is an artificial stimulus, our findings suggest that Mtb keeps in check its massive pro-inflammatory potential by the organization and integrity of the envelop (**Supplemental Fig. 6B**). We imagine that by altering cell envelop architecture, Mtb tune their interactions to achieve the desired host response (Garcia-Vilanova et al., 2019); for example, for initial infection and persistence, it may benefit the bacilli to minimize the TLR2-driven inflammatory response to promote immune evasion, whereas in order to drive tissue pathology and transmission, the bacilli may generate a hyperinflammatory phenotype (Chandra et al., 2022). While the Mtb cell wall is known to be dynamic (Dulberger et al., 2020), little is known about the structure and function of the cell wall during different *in vivo* contexts. To this end, a recent study evaluated the ultrastructure of the Mtb cell wall *ex vivo* from infected human sputum samples (Vijay et al., 2017). The characteristic three layers were found, and a reduction in the electron translucent layer was noted when bacilli were grown under stress conditions. Further *in vivo* studies investigating how Mtb regulates cell envelop architecture to modulate host inflammatory responses and deploy virulence lipids and protein effectors are needed.

## Materials and Methods

### Bacterial Strains and Growth Conditions

The Mtb strains H37Rv (WT), Δ*ppsD*, and Δ*ppsD∷ppsD* were used in this study. The Δ*ppsD* and Δ*ppsD∷ppsD* strains were from A. Barczak and previously described (Barczak et al., 2017). HN878 strain was a gift from Shabaana Khader, and Erdman was from Christina Stallings. Bacteria were grown to mid-log phase in an incubator at 37°C with 5% CO_2_ and gentle agitation (120 rpm). Bacteria were grown in 7H9 media supplemented with Middlebrook OADC (oleic acid, albumin, dextrose, catalase), 0.05% Tyloxapol, and 0.2% glycerol. H37Rv Δ*ppsD* growth media was additionally supplemented with 50 μg/mL hygromycin, GFP-expressing bacterial strains with 25 μg/mL kanamycin, and Δ*ppsD*∷*ppsD* with 50μg/mL hygromycin and 25μg/mL kanamycin.

### Generation of single cell suspensions of Mtb

Following growth of Mtb to mid-log phase (OD_600_ 0.5-0.8), bacteria were washed with phosphate-buffered saline (PBS) and resuspended in the appropriate media for the subsequent study. Single cell suspensions of Mtb were generated using one or a combination of the following methods: 1) low-speed spin (sp): bacteria were centrifuged at 206 × g for 10 min followed by 132 × g for 8 min, with the supernatant collected following each spin; 2) 5μm filter (5μmF): 6-20 ml of bacterial culture were added to a 10 mL syringe and then, with gentle pressure applied to the syringe plunger, passed through a 5μm polyethersulfone (PES) filter (PALL Life Sciences; cat. 4650) except in the case were polytetrafluoroethylene (PTFE) filters (Tisch scientific; cat. SF17400) were used; 3) sonication (so): 4-10 ml bacteria in a 15 mL conical tube were placed in a water bath sonicator (Branson Ultrasonics Corporation, Digital Sonifier 450) and sonicated with three pulses lasting 10 s each, with an amplitude of 70% and 5 s rests between each pulse. Following sonication, bacteria were centrifuged with a low-speed spin (so/sp) or passed through a 5μm filter (so/5μmF), as described above. Following the preparations described above, the concentrations of bacterial suspensions with OD_600_ between 0.04-0.12 were calculated using the formula: 1 OD_600_ = 3 × 10^8^ bacteria per mL (except filtered bacterial suspensions, which were calculated using: 1 OD_600_ = 2 × 10^8^ bacteria per mL). The multiplicity of infection was determined by plating the input on 7H10 plates. For some experiments, bacterial cultures were further passed through a 0.2 μm filter (PALL Life Sciences; cat. 4652).

### Mice

8- to 12-week-old C57BL/6J and *Tlr2*^−/−^ (B6.129-Tlr2tm1Kir/J) mice were obtained from The Jackson Laboratory. All work with mice were approved by the Washington University School of Medicine Institutional Animal Care and Use Committee. Euthanasia was performed prior to bone marrow harvest in accordance with the 2020 *AVMA Guidelines for the Euthanasia of Animals* prior to tissue harvest.

### Bone marrow-derived macrophage isolation and infection

Mouse hematopoietic stem cells were isolated as described in (Banaiee et al., 2006). Hematopoietic cells were differentiated by culturing for 7 days in Dulbecco’s Modified Eagle Medium (DMEM) with 10% FBS, 2 mM L-glutamine, and 1 mM pyruvate (DMEM complete). DMEM complete media was supplemented with 20% L929 cell supernatant (as a source of macrophage colony stimulating factor (M-CSF), 100 U/mL final concentration), 10 units/ml penicillin, and 10 units/ml streptomycin. Following differentiation, BMDMs were washed with PBS, resuspended in DMEM complete with 10% L929 cell supernatant, and plated for infection the following day. Single cell suspensions of Mtb in DMEM complete with 10% L929 added to macrophages at a MOI of 5, 10, 20, or 40, and plates were spun for 5 minutes at 51 × g. The MOI was verified by plating the inoculum. At 4 hpi, macrophages were washed 3 times with DMEM to remove extracellular bacteria. To enumerate CFU, at specified time points macrophages were lysed with 0.06% sodium dodecyl sulfate (SDS) in water and serially diluted in PBS. The cell lysates were plated on 7H11 agar plates supplemented with OADC and glycerol, and CFU were counted after 14-21 days. For qPCR, macrophages were lysed in TRI Reagent (Zymo Research, R2050-1-50), and total RNA was extracted.

### RNA sequencing

Mouse hematopoietic cells were collected and differentiated to BMDMs as above. 1.6 × 10^6^ BMDMs per well were incubated overnight in a six-well plate. BMDMs were either uninfected or infected at a MOI of 5 with Mtb prepared by the designated single cell preparation method. 5 samples per group were used. At 72 hpi, macrophages were lysed in TRI Reagent and total RNA was extracted. Total RNA integrity was determined using Agilent Bioanalyzer or 4200 Tapestation. Library preparation was performed with 500ng to 1ug total RNA. Ribosomal RNA was removed by an Rnase-H method using RiboErase kits (Kapa Biosystems). mRNA was then fragmented in reverse transcriptase buffer and heating to 94°C for 8 min. mRNA was reverse transcribed to yield cDNA using SuperScript III RT enzyme (Life Technologies, per manufacturer’s instructions) and random hexamers. A second strand reaction was performed to yield ds-cDNA. cDNA was blunt ended, had an A base added to the 3’ ends, and then had Illumina sequencing adapters ligated to the ends. Ligated fragments were then amplified for 12-15 cycles using primers incorporating unique dual index tags. Fragments were sequenced on an Illumina NovaSeq-6000 using paired end reads extending 150 bases. The raw CPM values that were generated underwent filtering, with removal of mitochondrial RNA, autosomal rRNA, and low-expressed genes with less than 1 CPM in the smallest group size, followed by Voom transformation of counts. Differentially expressed genes were then determined using the “limma” package from bioconductor.org. Heatmaps were generated in R using the “pheatmap” package.

### Gene set enrichment analysis

We imputed normalized gene expression data and associated Ensembl Stable IDs of differentially expressed genes from our RNA-seq experiment into GSEA software. GSEA then analyzed our dataset for enriched genetic signatures curated in the hallmark gene sets by the Molecular Signatures Database (Liberzon et al., 2015; Subramanian et al., 2005). Genes were ranked based on their expression and compared against the hallmark gene sets in order to generate an enrichment score. A nominal P value was then generated followed by normalization for the size of the gene set and adjustment for multiple hypothesis testing to yield a false discovery rate (FDR) as previously described (Subramanian et al., 2005). Gene sets which had a P-value <0.01 and an FDR <0.01 were considered significant.

### Quantitative Polymerase Chain Reaction

BMDMs (2.0 × 10^5^ per well) were plated in 24 well plates. At indicated time points, macrophage growth media was aspirated, and 100μL TRIzol (Zymo Research, R2050-1-50) was added to each well followed by isolation of total RNA using Direct-Zol RNA Mini-Prep Plus Kit (Zymo Research, R1058) according to the manufacturer’s instructions. RNA concentrations were determined using NanoDrop One (Thermo Fisher Scientific Inc., Waltham, MA), and cDNA was made with High Capacitance cDNA Reverse Transcription Kit (Thermo Fisher Scientific Inc., Waltham, MA). Quantitative PCR was performed using SYBR Green dye (CFX Connect Real-Time System, Bio-Rad Laboratories, Inc., Hercules, CA). Fold-changes in gene expression were calculating by normalizing data to *Gapdh* as a house-keeping gene and values were presented relative to uninfected cells. The nucleotide sequences of all primers used are presented in **Supplemental Table 2**.

### Macrophage viability assay

BMDMs were plated in 200μL of media in a 96-well white optical plate. After BMDMs were allowed to adhere, cells were infected with Mtb of the appropriate preparation at an MOI of 10. The plates were centrifuged at 51 × g for 5 minutes and incubated at 37°C and 5% CO_2_. At 4 hpi, macrophages were washed 3 times with DMEM to remove extracellular bacteria. At the appropriate time points, macrophage viability was determined using the CellTiter-Glo Luminescent Cell Viability Assay (Promega, catalog number G7570). At 4 hpi, 3 dpi, and 5 dpi, the media was aspirated and a solution of 100μL DMEM and 25μL CellTiter-Glo solution was added to each well per the manufacturer’s instructions. The plates were incubated at 37°C for 10 minutes, covered with optical tape, and luminescence was then determined using a Synergy HTX Multi-Mode Reader (Agilent Technologies, Inc., Santa Clara, CA). The relative luminescence units were normalized to the reading for uninfected BMDM samples at each given timepoint. Six biological replicates per group were performed at each timepoint.

### Fluorescence microscopy

BMDMs (3 × 10^4^ per well) were seeded in glass bottom 96-well plate (Ibidi, catalog number 89626) and infected with GFP-expressing H37Rv at a MOI of 5. After 4 h, macrophages were washed with PBS and fixed with 1% paraformaldehyde in PBS overnight followed by permeabilization in 0.1% vol/vol Triton X-100 (Millipore Sigma) in PBS for 10 min at room temperature (RT) and blocked for 45 min in 2% bovine serum albumin (BSA) in PBS prior to staining with DAPI (4=,6-diamidino-2-phenylindole) and mounted in Prolong Diamond antifade (Molecular Probes, Life Technologies). Images were captured using a Nikon Eclipse Ti confocal microscope (Nikon Instruments, Inc., Melville, NY) equipped with a 60X apochromat oil objective lens. Image acquisition was done using NIS-Elements version 4.40. Fluorescent images were then used to further analyze the bacterial clumps based upon manual quantification of GFP-bacteria in infected macrophages. Single bacteria and clumps of bacteria were quantified for each of the preparations and at least 100 bacterial occurrences were analyzed for each preparation method.

### ELISA

2.0 × 10^5^ BMDM in 24 well plates were infected with Mtb at a MOI of 10. At the indicated timepoints, the cell supernatant was collected and filtered through 0.22 μm filters. Cytokines were measured from the supernatant with R&D Systems DuoSet ELISA kits for TNF-α, IL1B, and IL6 according to the manufacturer’s instructions (R&D Systems, cat. DY406, DY410, DY411). Three biological replicates per group and two technical replicates per sample were used, and experiments were repeated at least two times per experimental condition.

### FluoroDOT assay

Assays were performed using reagents from Mouse TNF-α DuoSet ELISA kits (R&D systems, catalog number DY410-05). Wells of 96 well glass-bottom, black plates (P96-1.5H-N, Cellvis, Mountain View, USA) were coated with 100μL TNF-α capture antibody (2 μg/mL in PBS) at 4°C overnight. Coated wells were then washed 3 times with PBS, followed by blocking with reagent diluent (0.2μm filtered 1% BSA in PBS) for at least 1 h at RT. Wells were washed 3 times with PBS and thereafter 8.0 × 10^3^ BMDMs in DMEM complete with 10% L cell supernatant were added to each well. The same day, BMDMs were infected with the indicated GFP-expressing Mtb strains that had prepared as single cell suspensions. Macrophages were incubated at 37°C in 5% CO_2_ for 3 h, followed by 3 washes with fresh media to remove extracellular Mtb, and incubated in the same media for an additional 3 h. Media was then aspirated and 200μL 4% PFA in PBS was added for 30 min at 37°C. Wells were washed with PBS and incubated with biotinylated 75 ng/mL TNF-α detection antibody in reagent diluent for 2 h at RT. Wells were washed 3 times with PBS followed by 100μL PBS containing streptavidin plasmonic-fluor 650 (PF650, extinction 0.5; Auragent Bioscience LLC) (Wang et al., 2021) for 30 min at RT in the dark. Cells were washed 3 times with PBS and stained with 300nM DAPI (Millipore Sigma) for 5 min at RT in the dark. Wells were washed 3 times with PBS and then visualized using a Nikon TsR2 epifluorescence microscope.

### Transmission electron microscopy

Bacteria were grown to mid-log phase, and single cell suspensions were generated in PBS as described above. Bacteria were incubated in 4% PFA for 30 min at 37°C followed by centrifugation at 3000 × g and resuspension in PBS. For ultrastructural analyses using ultrathin cross-sections through bacteria, samples were further fixed in 2% paraformaldehyde/2.5% glutaraldehyde (Ted Pella Inc., Redding, CA) in 100 mM sodium cacodylate buffer, pH 7.2 for 2h at RT and then overnight at 4°C. Samples were washed in sodium cacodylate buffer at RT and postfixed in 2% osmium tetroxide (Ted Pella Inc) for 1h at RT. Samples were then rinsed in dH20, dehydrated in a graded series of ethanol, and embedded in Eponate 12 resin (Ted Pella Inc). Sections of 95 nm were cut with a Leica Ultracut UCT ultramicrotome (Leica Microsystems Inc., Bannockburn, IL), stained with uranyl acetate and lead citrate, and viewed on a JEOL 1200 EX transmission electron microscope (JEOL USA Inc., Peabody, MA) equipped with an AMT 8-megapixel digital camera and AMT Image Capture Engine V602 software (Advanced Microscopy Techniques, Woburn, MA).

For imaging of whole bacteria, after bacterial samples were fixed with 4% PFA, they were allowed to adsorb onto freshly glow discharged formvar/carbon-coated copper grids for 10 min. Grids were then washed in dH2O and stained with 1% aqueous uranyl acetate (Ted Pella Inc., Redding, CA) for 1 min. Excess liquid was gently wicked off, and grids were allowed to air dry. Samples were viewed by transmission electron microscopy as described above. TEM images were then used to further analyze the bacteria using ImageJ (version 1.53q). Single bacteria and clumps of bacteria which were completely contained within the image borders were further analyzed for each of the preparations. For each single bacteria or clump, a measurement was then taken of the thickness of the capsular layer. Lines were drawn perpendicular to the middle of the long axis of the bacteria, capturing the black-staining outer component.

### Methodology of literature review

We conducted a search in PubMed using the medical subject headings “*Mycobacterium tuberculosis*” and “Macrophage” and filtered for articles published in 2021. This generated 183 articles, which were further filtered to include only original research articles by removing review articles, protocols, and commentaries. The remaining 155 articles were included in the analysis if they performed an *in vitro* macrophage infection with live Mtb. The text, supplementary methods, and figures were reviewed to determine the single cell preparation methods used.

### Statistical analysis

Graph Pad Prism 9 software was used for statistical analysis and to prepare graphs. Error bars used in the figures correspond to the mean and standard deviation. Statistical significance was determined using unpaired T test, one-way analysis of variance (ANOVA), or two-way ANOVA, as indicated.

## Acknowledgements

We thank members of the Philips laboratory for their input. Funding for these studies came from NIAID/NIH (R01 AI087682 and AI30454) to JAP, and the National Cancer Institute (NCI)-Innovative Molecular Analysis Technologies (R21CA236652) and National Science Foundation (CBET-1900277) to SS. ATR was supported by NIH/NHLBI (T32 HL007317-37). We thank the Genome Technology Access Center at Washington University School of Medicine for help with genomic analysis. The Center is partially supported by NCI Cancer Center Support Grant #P30 CA91842 to the Siteman Cancer Center and by ICTS/CTSA Grant# UL1TR002345 from the National Center for Research Resources (NCRR), a component of the National Institutes of Health (NIH), and NIH Roadmap for Medical Research. This publication is solely the responsibility of the authors and does not necessarily represent the official view of NCRR or NIH.

## Data availability

RNA-seq data can be accessed in the Gene Expression Omnibus database (Accession GSE206485; ID: 200206485).

## Competing interests

SS is an inventor on a provisional patent related to plasmonic-fluor technology, and the technology has been licensed by the Office of Technology Management at Washington University in St. Louis to Auragent Bioscience LLC. SS is a co-founder/shareholder of Auragent Bioscience LLC. SS along with Washington University may have financial gain through Auragent Bioscience LLC through this licensing agreement. AS is currently employed with Auragent Bioscience LLC. These potential conflicts of interest have been disclosed and are being managed by Washington University in St. Louis.

## Author contributions

The project was conceptualized by EM and JAP. All experiments were performed by EM and ATR, and formal analysis was performed by ATR, EM, and JAP. AS performed FluroDOT experiments. SS supervised FluroDOT studies. WB performed electron microscopy experiments. Funding was acquired by JAP and SS. Writing of the original draft was done by ATR, EM, and JAP. Editing of the manuscript was done by all authors. JAP supervised the studies. EM and ATR share the first author position, reflecting their equivalent contributions.

## Supplementary Files

**Supplemental Table 1. RNA-seq data**

This file lists the genes that were differentially expressed between uninfected macrophages, macrophages infected with Mtb prepared by sonication followed by low-speed spin (so/sp), and macrophages infected with Mtb prepared by passing through a 5μm filter (5μmF). Infectious were carried out at an MOI of 5 and analyzed at 72 hpi.

**Supplemental Table 2.**
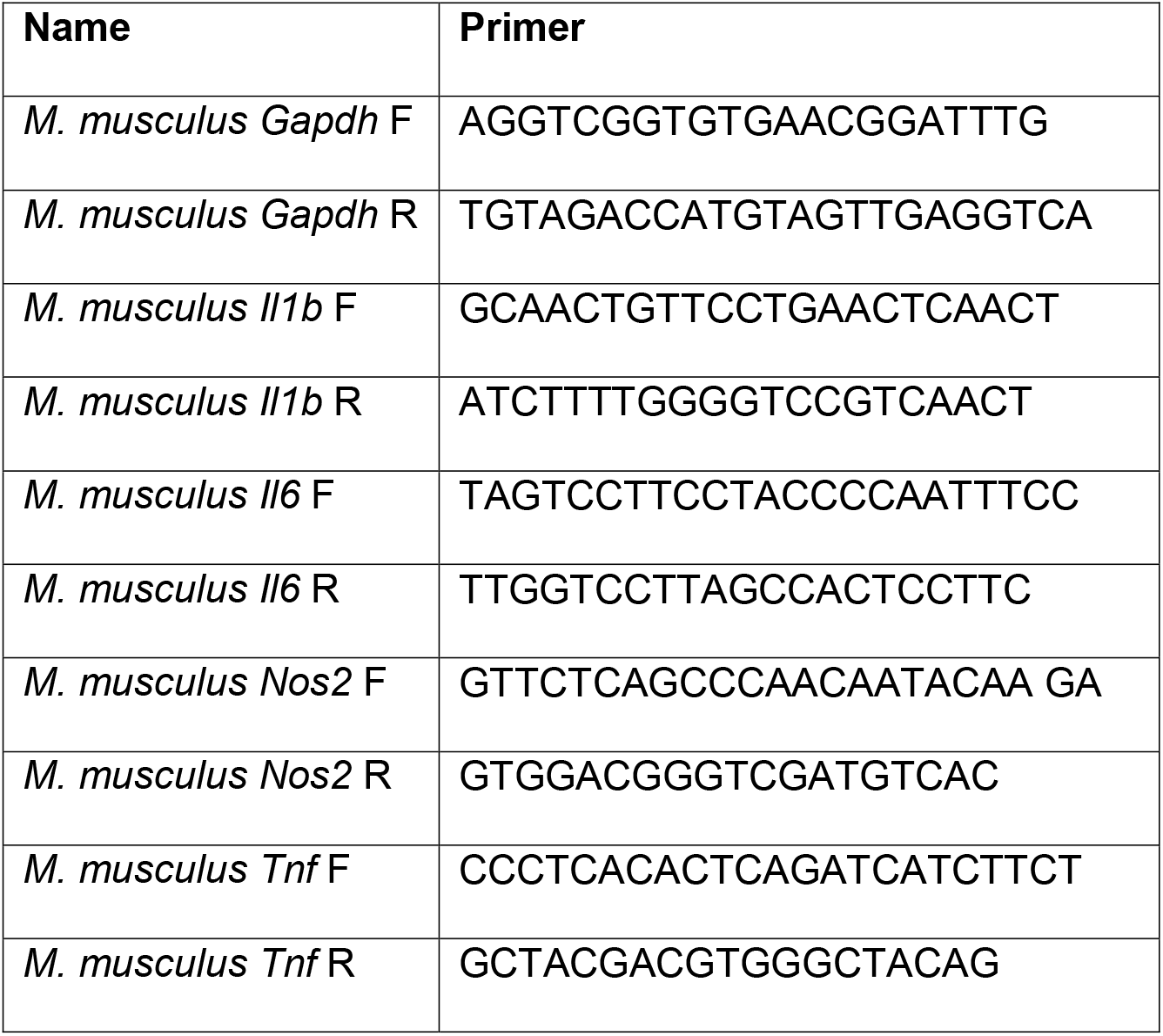
PCR primers used.

---

**Supplemental Figure 1.**
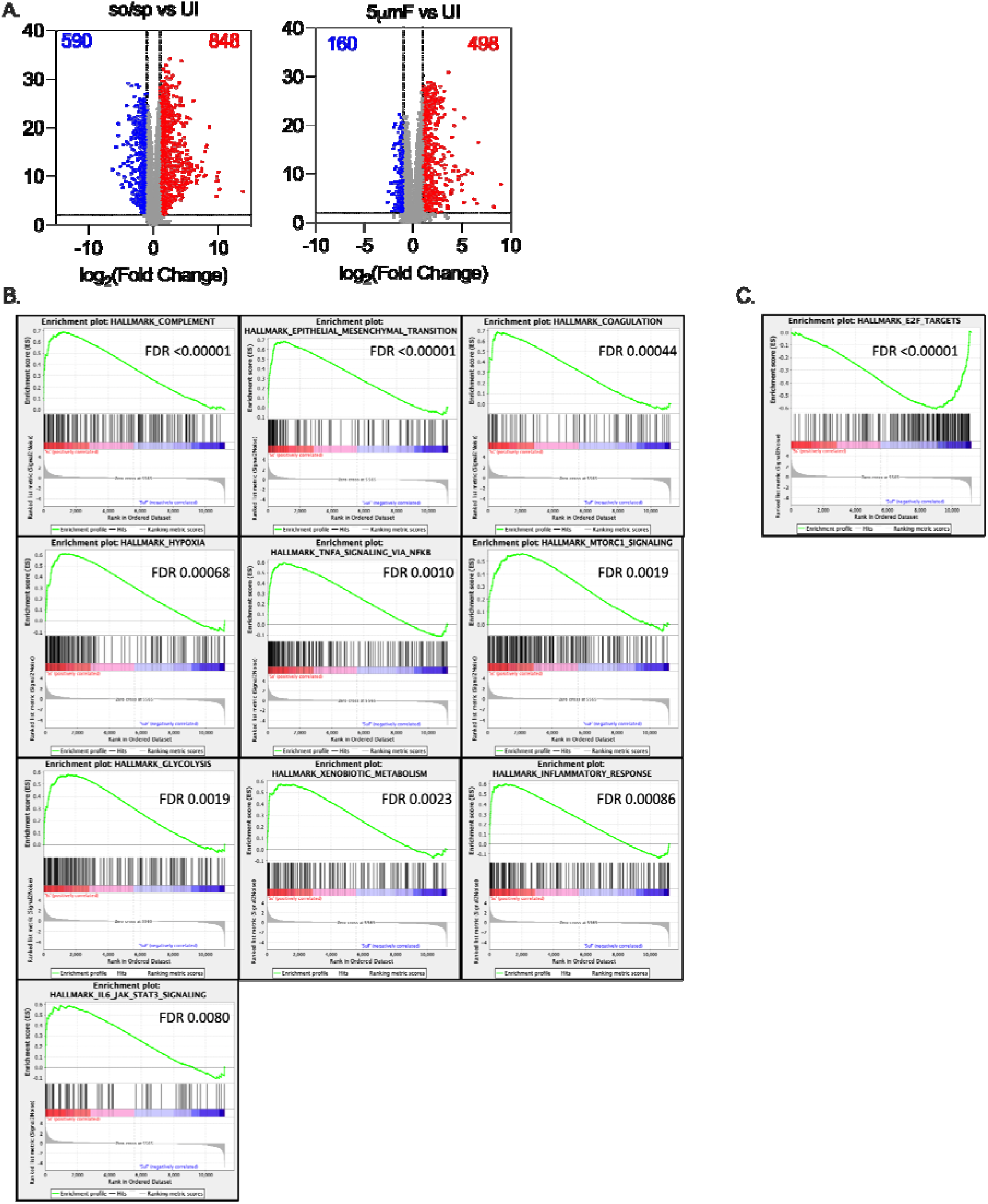
Infection-induced changes in macrophage gene expression depend on whether Mtb are sonicated or filtered. **(A)** Volcano plot shows genes differentially expressed in BMDMs infected with so/sp versus UI (left) or 5μmF Mtb versus UI (right). DEGs exhibiting an adjusted P-value of ≤0.01 and a linear fold change ≥2.00 (red) or ≤-2.00 (blue) are indicated. **(B-C)** Gene set enrichment analysis (GSEA) leading edge graphs of hallmark gene sets that were enriched in BMDMs infected with so/sp relative to 5μmF Mtb (B) or 5μmF relative to so/sp **(C)**.

**Supplemental Figure 2.**
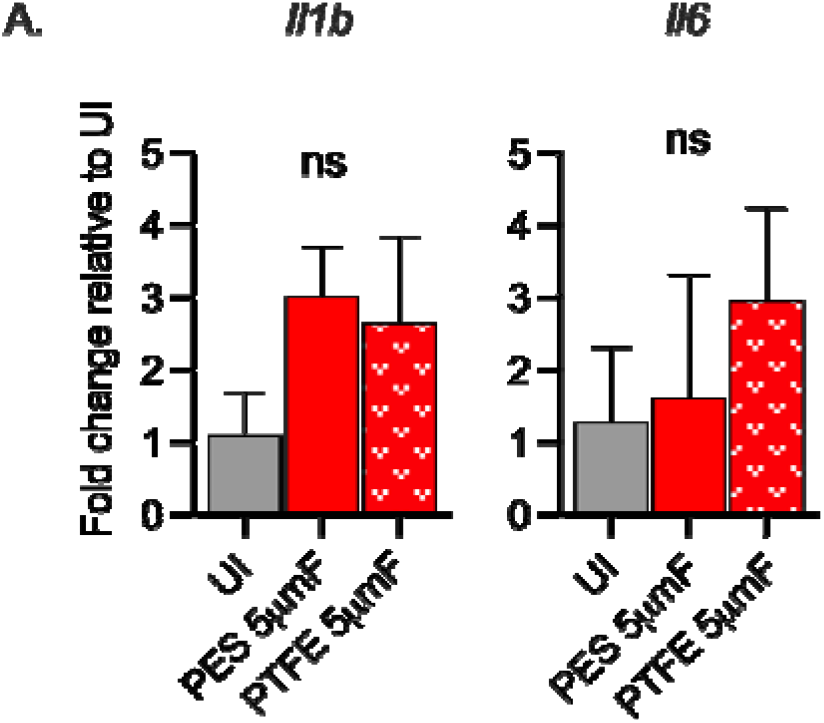
Filtered Mtb are uninflammatory irrespective of filter type. BMDMs were uninfected or infected with Mtb prepared by passing through a 5μmF made of either PES or PTFE using a MOI of 10 and analyzed by qPCR 6 hpi. Data are presented as fold changes in gene expression relative to uninfected BMDMs. Data are representative of 2 to 3 experiments, each with three biological replicates per group and two technical replicates per sample. Statistical significance was determined with one-way ANOVA using Tukey’s multiple comparisons test. qPCR data are presented as fold change in gene expression relative to uninfected BMDMs. Error bars indicate mean +/− SD. ns not significant.

**Supplemental Figure 3.**
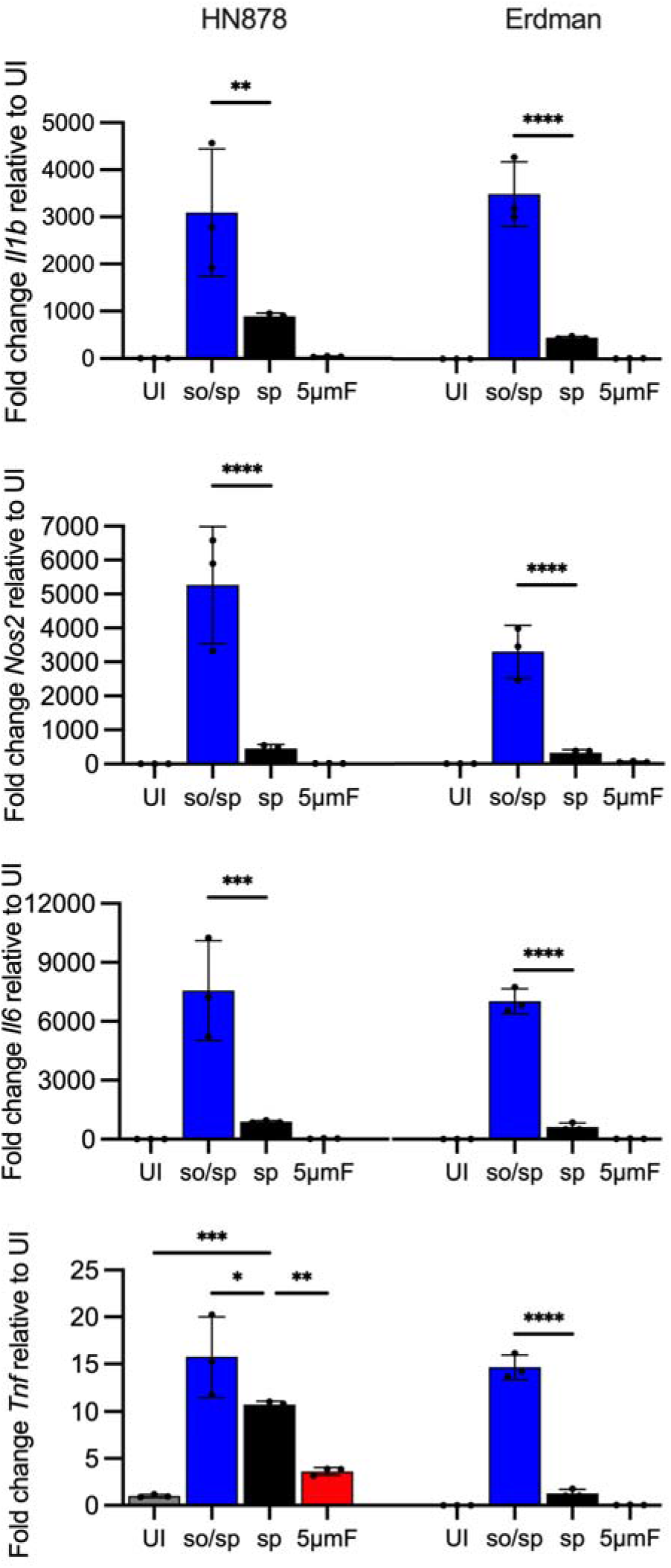
Sonicated HN878 and Erdman strains induce high TLR2-dependent inflammatory responses. BMDMs were uninfected (UI) or infected with different preparations of HN878 or Erdman as indicated at an MOI of 10 and analyzed by qPCR 6 hpi. Data are presented as fold changes in gene expression relative to uninfected BMDMs. Three biological replicates were used per group and two technical replicates per sample. Statistical significance was determined with one-way ANOVA using Tukey’s multiple comparisons test. Error bars indicate mean +/− SD. ns not significant; *P<0.05; **P<0.01; ***<0.001; ****P<0.0001.

**Supplemental Figure 4.**
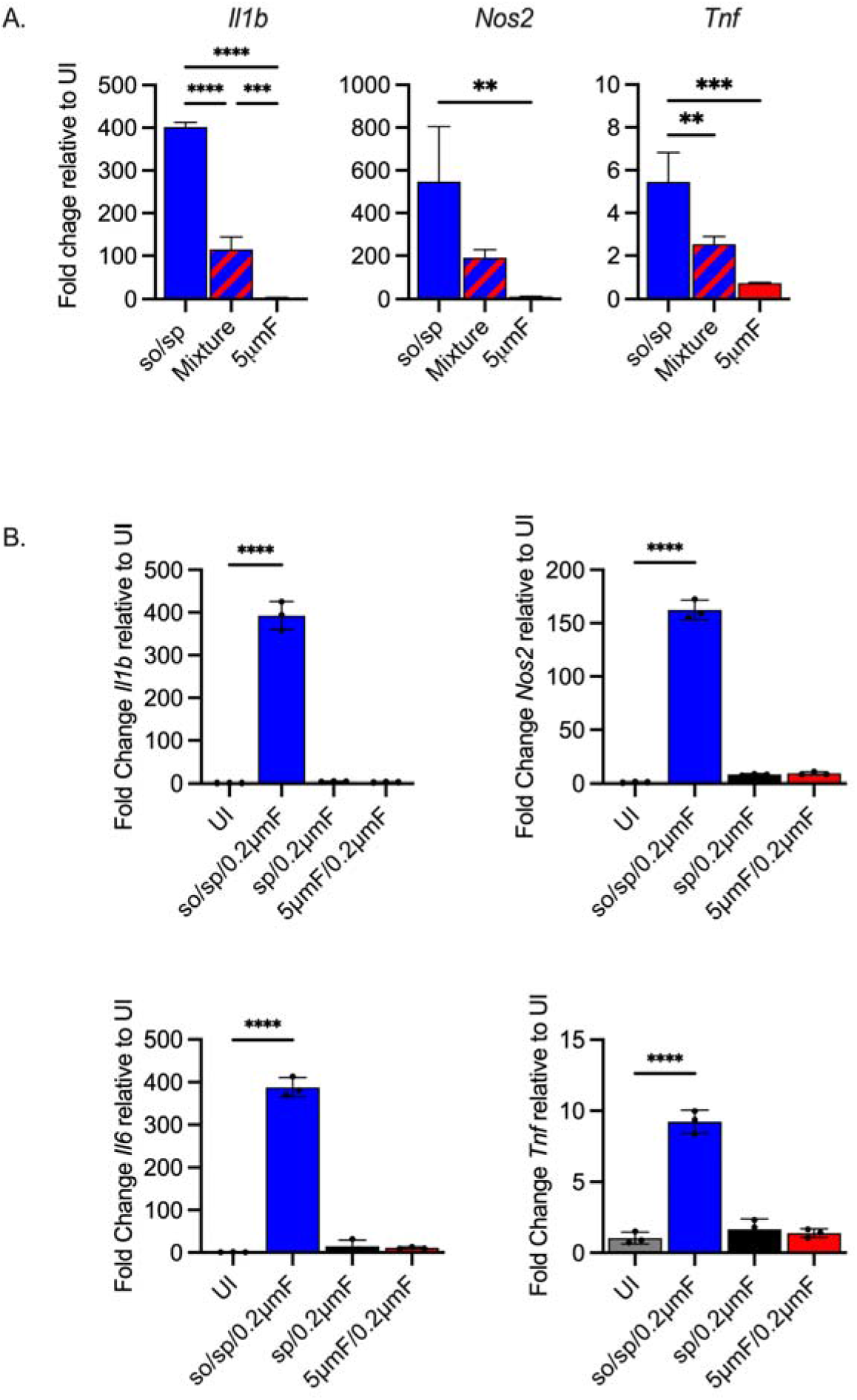
Filtered bacteria do not strongly inhibit response from sonicated bacteria and only the sterile filtrate of so/sp Mtb induces gene expression in BMDMs. **(A)** BMDMs were uninfected or infected for 6h at a MOI of 10 with bacteria prepared by the so/sp or 5μmF-preparation or a mixture (1:1) of the two samples. Data are presented as fold change in gene expression relative to uninfected BMDMs. Statistical significance was determined with one-way ANOVA using Tukey’s multiple comparisons test. **(B)** BMDMs were untreated, or treated with the sterile filtrate from different preparations for 6h and analyzed by qPCR. Data are presented as fold changes in gene expression relative to untreated BMDMs. Statistical significance was determined for each group relative to untreated BMDMs with one-way ANOVA using Dunnett’s multiple comparisons test. **(A-B)** Data are representative of 3 experiments, each with three biological replicates per group and two technical replicates per sample. Error bars indicate mean +/− SD. ns not significant; **P<0.01; ***<0.001; ****<0.0001.

**Supplemental Figure 5.**
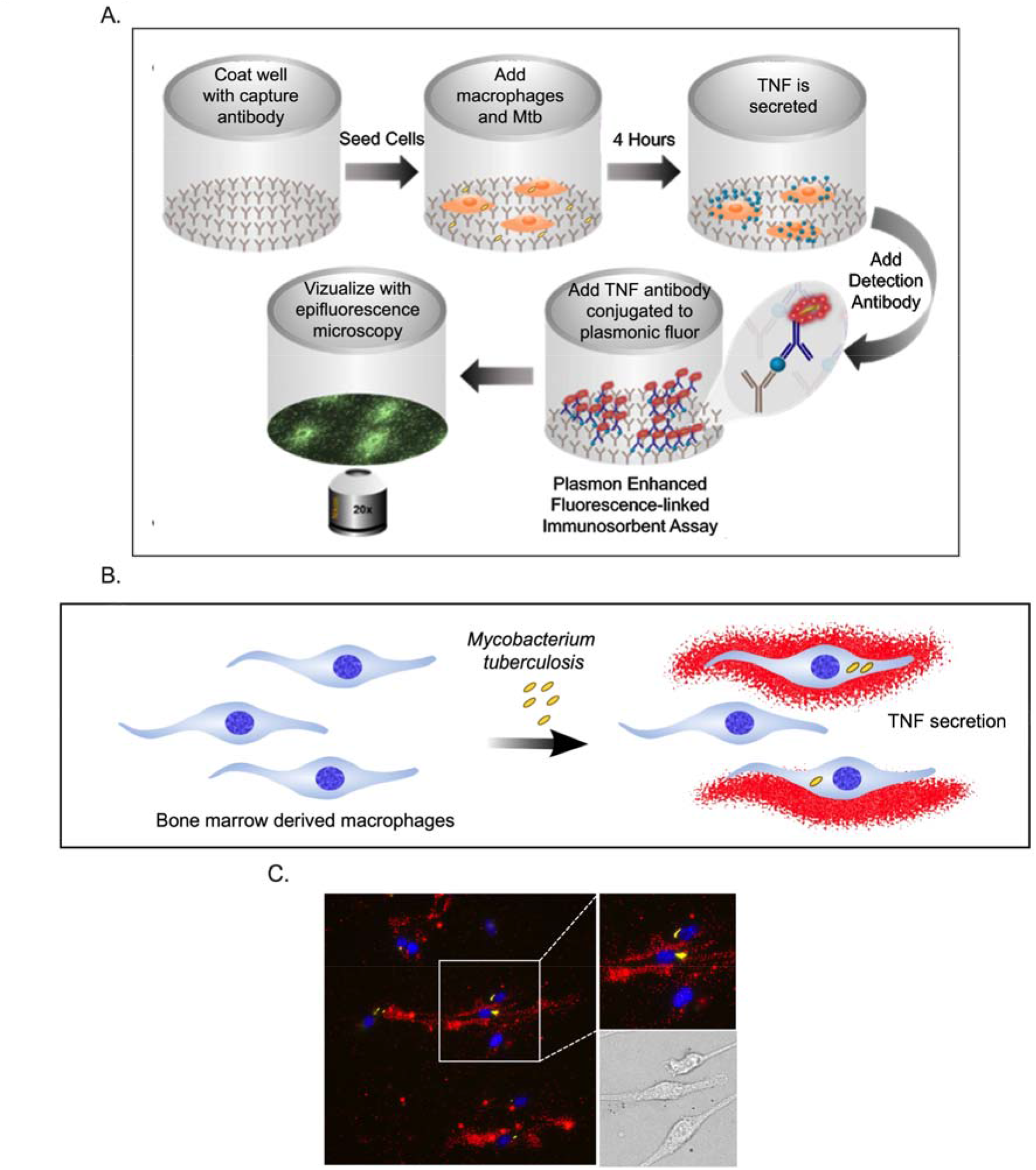
Schematic representation of the FluoroDOT assay. **(A)** 96-well glass bottom plates are coated with TNF-α capture antibody, followed by the addition of BMDMs. The macrophages are then infected with Mtb. TNF-α that is secreted by the BMDMs can be bound by the capture antibody. Samples are fixed and then the detection antibody, which is conjugated to plasmon-fluor 650, is added and the plate is visualized using epifluorescence microscopy**. (B)** Illustration indicating how the assay can reveal TNF-α secretion from Mtb-infected or bystander macrophages. **(C)** Representative fluorescent image with zoomed region and corresponding brightfield image. The three cells in the images demonstrate examples of an infected macrophage secreting substantial TNF-α (middle), an infected cell with little TNF-α secretion (top), and an uninfected within minimal TNF-α secretion (bottom). Macrophages were stained with DAPI (blue); TNF-α is red.

**Supplementary Figure 6.**
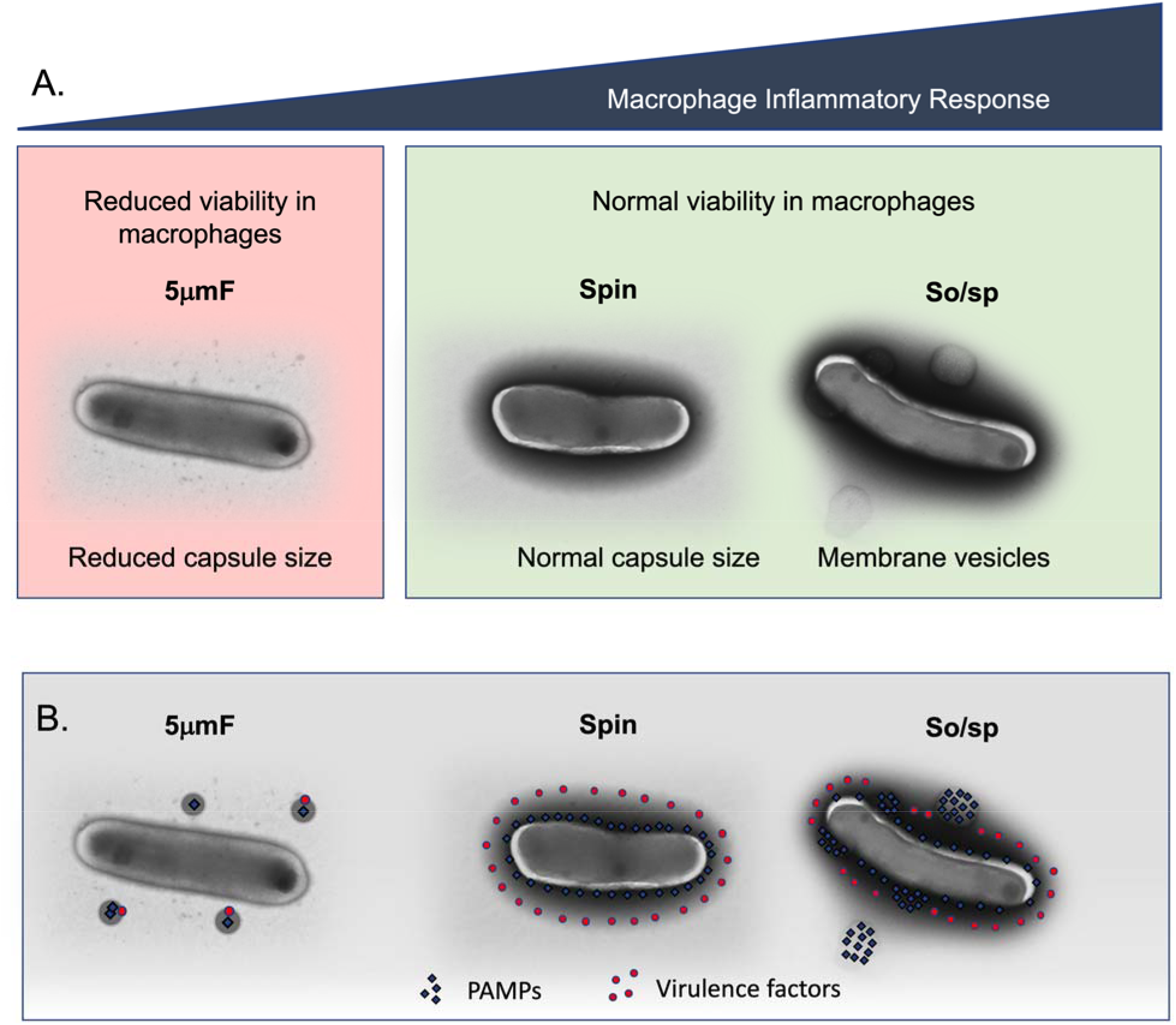
Summary of findings and model describing the impact of bacterial preparation methods on host-pathogen interactions. **(A)** Filtered (5μmF), spun (sp), and sonicated (so/sp) Mtb differ in appearance and elicit different macrophage responses. 5μmF bacteria exhibit reduced growth in macrophages and have reduced capsular staining. Bacilli prepared with sonication elicit the strongest inflammatory response and have membrane vesicles and cell envelop protrusions. **(B)** The findings in **(A)** can be explained by the following model: filtration disrupts the cell envelope, dispersing and inactivating PAMPs and virulence factors, resulting in reduced macrophage inflammatory responses and reduced intracellular growth. Bacteria that are spun have an intact cell envelop that shields PAMPs and contains virulence factors. Sonication disrupts the cell envelop such that PAMPs are more highly exposed, resulting in increased inflammatory responses; virulence factors remain intact enabling normal growth in macrophages.

**Supplemental Figure 7.**
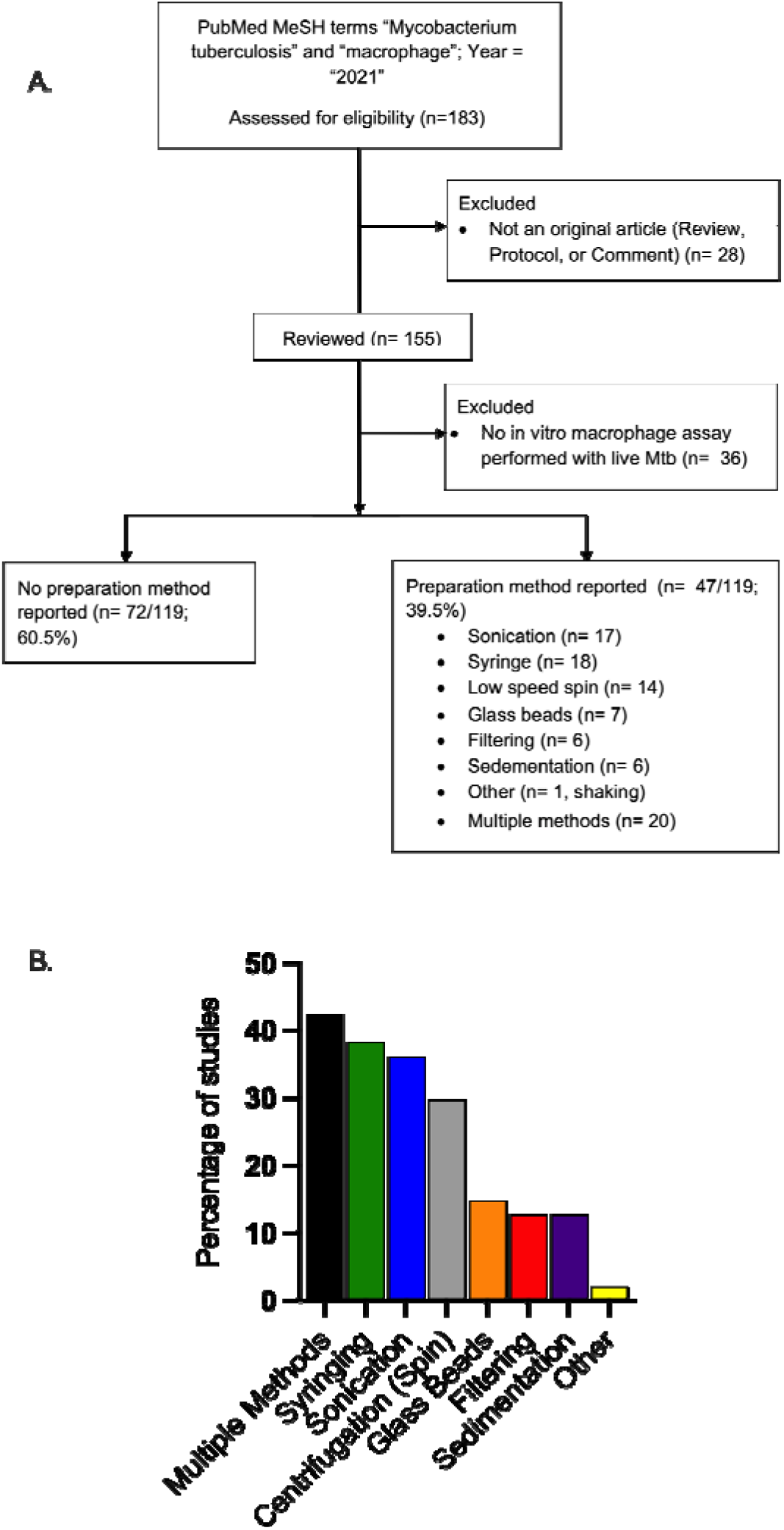
Literature review of methods used to generate single cell suspensions. **(A)** Approach used to analyze the literature to define the frequency with which distinct single cell preparation methods are used and how often they are reported. **(B)** Graph demonstrates the distribution of methods reported. Since some studies used multiple methods, the total does not equal 100.

## Notes

### Summary of Updates

There are a number of revisions that have been made, including incorporation of data that the Erdman and HN878 strains behave similarly to H37Rv.

## References

Astarie-Dequeker, C., Le Guyader, L., Malaga, W., Seaphanh, F. K., Chalut, C., Lopez, A., & Guilhot, C. (2009). Phthiocerol dimycocerosates of M. tuberculosis participate in macrophage invasion by inducing changes in the organization of plasma membrane lipids. PLoS Pathog, 5(2), e1000289. https://doi.org/10.1371/journal.ppat.1000289

Athman, J. J., Wang, Y., McDonald, D. J., Boom, W. H., Harding, C. V., & Wearsch, P. A. (2015). Bacterial Membrane Vesicles Mediate the Release of Mycobacterium tuberculosis Lipoglycans and Lipoproteins from Infected Macrophages. J Immunol, 195(3), 1044–1053. https://doi.org/10.4049/jimmunol.1402894

Augenstreich, J., Arbues, A., Simeone, R., Haanappel, E., Wegener, A., Sayes, F., … Astarie-Dequeker, C. (2017). ESX-1 and phthiocerol dimycocerosates of Mycobacterium tuberculosis act in concert to cause phagosomal rupture and host cell apoptosis. Cell Microbiol, 19(7). https://doi.org/10.1111/cmi.12726

Banaiee, N., Kincaid, E. Z., Buchwald, U., Jacobs, W. R., Jr., & Ernst, J. D. (2006). Potent inhibition of macrophage responses to IFN-gamma by live virulent Mycobacterium tuberculosis is independent of mature mycobacterial lipoproteins but dependent on TLR2. J Immunol, 176(5), 3019–3027. https://doi.org/10.4049/jimmunol.176.5.3019

Barczak, A. K., Avraham, R., Singh, S., Luo, S. S., Zhang, W. R., Bray, M. A., … Hung, D. T. (2017). Systematic, multiparametric analysis of Mycobacterium tuberculosis intracellular infection offers insight into coordinated virulence. PLoS Pathog, 13(5), e1006363. https://doi.org/10.1371/journal.ppat.1006363

Blanc, L., Gilleron, M., Prandi, J., Song, O. R., Jang, M. S., Gicquel, B., … Nigou, J. (2017). Mycobacterium tuberculosis inhibits human innate immune responses via the production of TLR2 antagonist glycolipids. Proc Natl Acad Sci U S A, 114(42), 11205–11210. https://doi.org/10.1073/pnas.1707840114

Camacho, L. R., Constant, P., Raynaud, C., Laneelle, M. A., Triccas, J. A., Gicquel, B., … Guilhot, C. (2001). Analysis of the phthiocerol dimycocerosate locus of Mycobacterium tuberculosis. Evidence that this lipid is involved in the cell wall permeability barrier. J Biol Chem, 276(23), 19845–19854. https://doi.org/10.1074/jbc.M100662200

Cambier, C. J., Banik, S. M., Buonomo, J. A., & Bertozzi, C. R. (2020). Spreading of a mycobacterial cell-surface lipid into host epithelial membranes promotes infectivity. Elife, 9. https://doi.org/10.7554/eLife.60648

Cambier, C. J., Takaki, K. K., Larson, R. P., Hernandez, R. E., Tobin, D. M., Urdahl, K. B., … Ramakrishnan, L. (2014). Mycobacteria manipulate macrophage recruitment through coordinated use of membrane lipids. Nature, 505(7482), 218–222. https://doi.org/10.1038/nature12799

Chandra, P., Grigsby, S. J., & Philips, J. A. (2022). Immune evasion and provocation by Mycobacterium tuberculosis. Nat Rev Microbiol. https://doi.org/10.1038/s41579-022-00763-4

Cheng, N., Porter, M. A., Frick, L. W., Nguyen, Y., Hayden, J. D., Young, E. F., … Janzen, W. P. (2014). Filtration improves the performance of a high-throughput screen for anti-mycobacterial compounds. PLoS One, 9(5), e96348. https://doi.org/10.1371/journal.pone.0096348

Cox, J. S., Chen, B., McNeil, M., & Jacobs, W. R., Jr. (1999). Complex lipid determines tissue-specific replication of Mycobacterium tuberculosis in mice. Nature, 402(6757), 79–83.

Dinkele, R., Gessner, S., McKerry, A., Leonard, B., Seldon, R., Koch, A. S., … Warner, D. F. (2021). Capture and visualization of live Mycobacterium tuberculosis bacilli from tuberculosis patient bioaerosols. PLoS Pathog, 17(2), e1009262. https://doi.org/10.1371/journal.ppat.1009262

Dulberger, C. L., Rubin, E. J., & Boutte, C. C. (2020). The mycobacterial cell envelope - a moving target. Nat Rev Microbiol, 18(1), 47–59. https://doi.org/10.1038/s41579-019-0273-7

Garcia-Vilanova, A., Chan, J., & Torrelles, J. B. (2019). Underestimated Manipulative Roles of M*ycobacterium tuberculosis* Cell Envelope Glycolipids During Infection. Front Immunol, 10, 2909. https://doi.org/10.3389/fimmu.2019.02909

Hinman, A. E., Jani, C., Pringle, S. C., Zhang, W. R., Jain, N., Martinot, A. J., & Barczak, A. K. (2021). Mycobacterium tuberculosis canonical virulence factors interfere with a late component of the TLR2 response. Elife, 10. https://doi.org/10.7554/eLife.73984

Kalscheuer, R., Palacios, A., Anso, I., Cifuente, J., Anguita, J., Jacobs, W. R., … Prados-Rosales, R. (2019). The *Mycobacterium tuberculosis* capsule: a cell structure with key implications in pathogenesis. Biochem J, 476(14), 1995–2016. https://doi.org/10.1042/BCJ20190324

Kolloli, A., Kumar, R., Singh, P., Narang, A., Kaplan, G., Sigal, A., & Subbian, S. (2021). Aggregation state of Mycobacterium tuberculosis impacts host immunity and augments pulmonary disease pathology. Commun Biol, 4(1), 1256. https://doi.org/10.1038/s42003-021-02769-9

Lemassu, A., & Daffe, M. (1994). Structural features of the exocellular polysaccharides of Mycobacterium tuberculosis. Biochem J, 297 (Pt 2), 351–357. https://doi.org/10.1042/bj2970351

Lemassu, A., Ortalo-Magne, A., Bardou, F., Silve, G., Laneelle, M. A., & Daffe, M. (1996). Extracellular and surface-exposed polysaccharides of non-tuberculous mycobacteria. Microbiology (Reading), 142 (Pt 6), 1513–1520. https://doi.org/10.1099/13500872-142-6-1513

Lerner, T. R., Queval, C. J., Fearns, A., Repnik, U., Griffiths, G., & Gutierrez, M. G. (2018). Phthiocerol dimycocerosates promote access to the cytosol and intracellular burden of Mycobacterium tuberculosis in lymphatic endothelial cells. BMC Biol, 16(1), 1. https://doi.org/10.1186/s12915-017-0471-6

Liberzon, A., Birger, C., Thorvaldsdottir, H., Ghandi, M., Mesirov, J. P., & Tamayo, P. (2015). The Molecular Signatures Database (MSigDB) hallmark gene set collection. Cell Syst, 1(6), 417–425. https://doi.org/10.1016/j.cels.2015.12.004

Mahamed, D., Boulle, M., Ganga, Y., Mc Arthur, C., Skroch, S., Oom, L., … Sigal, A. (2017). Intracellular growth of Mycobacterium tuberculosis after macrophage cell death leads to serial killing of host cells. Elife, 6. https://doi.org/10.7554/eLife.22028

Murry, J. P., Pandey, A. K., Sassetti, C. M., & Rubin, E. J. (2009). Phthiocerol dimycocerosate transport is required for resisting interferon-gamma-independent immunity. J Infect Dis, 200(5), 774–782. https://doi.org/10.1086/605128

Ortalo-Magne, A., Dupont, M. A., Lemassu, A., Andersen, A. B., Gounon, P., & Daffe, M. (1995). Molecular composition of the outermost capsular material of the tubercle bacillus. Microbiology (Reading), 141 (Pt 7), 1609–1620. https://doi.org/10.1099/13500872-141-7-1609

Osman, M. M., Pagán, A. J., Shanahan, J. K., & Ramakrishnan, L. (2020). Mycobacterium marinum phthiocerol dimycocerosates enhance macrophage phagosomal permeabilization and membrane damage. PLoS One, 15(7), e0233252. https://doi.org/10.1371/journal.pone.0233252

Palacios, A., Gupta, S., Rodriguez, G. M., & Prados-Rosales, R. (2021). Extracellular vesicles in the context of Mycobacterium tuberculosis infection. Mol Immunol, 133, 175–181. https://doi.org/10.1016/j.molimm.2021.02.010

Prados-Rosales, R., Baena, A., Martinez, L. R., Luque-Garcia, J., Kalscheuer, R., Veeraraghavan, U., … Casadevall, A. (2011). Mycobacteria release active membrane vesicles that modulate immune responses in a TLR2-dependent manner in mice. J Clin Invest, 121(4), 1471–1483. https://doi.org/10.1172/JCI44261

Prados-Rosales, R., Carreno, L. J., Weinrick, B., Batista-Gonzalez, A., Glatman-Freedman, A., Xu, J., … Casadevall, A. (2016). The Type of Growth Medium Affects the Presence of a Mycobacterial Capsule and Is Associated With Differences in Protective Efficacy of BCG Vaccination Against Mycobacterium tuberculosis. J Infect Dis, 214(3), 426–437. https://doi.org/10.1093/infdis/jiw153

Quigley, J., Hughitt, V. K., Velikovsky, C. A., Mariuzza, R. A., El-Sayed, N. M., & Briken, V. (2017). The Cell Wall Lipid PDIM Contributes to Phagosomal Escape and Host Cell Exit of Mycobacterium tuberculosis. mBio, 8(2).https://doi.org/10.1128/mBio.00148-17

Reed, M. B., Domenech, P., Manca, C., Su, H., Barczak, A. K., Kreiswirth, B. N., … Barry, C. E. (2004). A glycolipid of hypervirulent tuberculosis strains that inhibits the innate immune response. Nature, 431(7004), 84–87. https://doi.org/10.1038/nature02837

Rodel, H. E., Ferreira, I., Ziegler, C. G. K., Ganga, Y., Bernstein, M., Hwa, S. H., … Sigal, A. (2021). Aggregated Mycobacterium tuberculosis Enhances the Inflammatory Response. Front Microbiol, 12, 757134. https://doi.org/10.3389/fmicb.2021.757134

Rousseau, C., Winter, N., Pivert, E., Bordat, Y., Neyrolles, O., Ave, P., … Jackson, M. (2004). Production of phthiocerol dimycocerosates protects Mycobacterium tuberculosis from the cidal activity of reactive nitrogen intermediates produced by macrophages and modulates the early immune response to infection. Cell Microbiol, 6(3), 277–287. https://doi.org/10.1046/j.1462-5822.2004.00368.x

Sani, M., Houben, E. N., Geurtsen, J., Pierson, J., de Punder, K., van Zon, M., … Peters, P. J. (2010). Direct visualization by cryo-EM of the mycobacterial capsular layer: a labile structure containing ESX-1-secreted proteins. PLoS Pathog, 6(3), e1000794. https://doi.org/10.1371/journal.ppat.1000794

Seth, A., Mittal, E., Luan, J., Kolla, S., Mazer, M. B., Joshi, H., … Singamaneni, S. (2022). High-resolution imaging of protein secretion at the single-cell level using plasmon-enhanced FluoroDOT assay. Cell Reports Methods, 2(8), 100267. https://doi.org/https://doi.org/10.1016/j.crmeth.2022.100267

Siméone, R., Constant, P., Malaga, W., Guilhot, C., Daffé, M., & Chalut, C. (2007). Molecular dissection of the biosynthetic relationship between phthiocerol and phthiodiolone dimycocerosates and their critical role in the virulence and permeability of Mycobacterium tuberculosis. FEBS J, 274(8), 1957–1969. https://doi.org/10.1111/j.1742-4658.2007.05740.x

Stokes, R. W., Norris-Jones, R., Brooks, D. E., Beveridge, T. J., Doxsee, D., & Thorson, L. M. (2004). The glycan-rich outer layer of the cell wall of Mycobacterium tuberculosis acts as an antiphagocytic capsule limiting the association of the bacterium with macrophages. Infect Immun, 72(10), 5676–5686. https://doi.org/10.1128/IAI.72.10.5676-5686.2004

Stukalov, O., Korenevsky, A., Beveridge, T. J., & Dutcher, J. R. (2008). Use of atomic force microscopy and transmission electron microscopy for correlative studies of bacterial capsules. Appl Environ Microbiol, 74(17), 5457–5465. https://doi.org/10.1128/AEM.02075-07

Subramanian, A., Tamayo, P., Mootha, V. K., Mukherjee, S., Ebert, B. L., Gillette, M. A., … Mesirov, J. P. (2005). Gene set enrichment analysis: a knowledge-based approach for interpreting genome-wide expression profiles. Proc Natl Acad Sci U S A, 102(43), 15545–15550. https://doi.org/10.1073/pnas.0506580102

Ufimtseva, E. G., Eremeeva, N. I., Petrunina, E. M., Umpeleva, T. V., Bayborodin, S. I., Vakhrusheva, D. V., & Skornyakov, S. N. (2018). Mycobacterium tuberculosis cording in alveolar macrophages of patients with pulmonary tuberculosis is likely associated with increased mycobacterial virulence. Tuberculosis (Edinb), 112, 1–10. https://doi.org/10.1016/j.tube.2018.07.001

Vijay, S., Hai, H. T., Thu, D. D. A., Johnson, E., Pielach, A., Phu, N. H., … Thuong, N. T. T. (2017). Ultrastructural Analysis of Cell Envelope and Accumulation of Lipid Inclusions in Clinical Mycobacterium tuberculosis Isolates from Sputum, Oxidative Stress, and Iron Deficiency. Front Microbiol, 8, 2681. https://doi.org/10.3389/fmicb.2017.02681

Wang, Z., Luan, J., Seth, A., Liu, L., You, M., Gupta, P., … Singamaneni, S. (2021). Microneedle patch for the ultrasensitive quantification of protein biomarkers in interstitial fluid. Nat Biomed Eng, 5(1), 64–76. https://doi.org/10.1038/s41551-020-00672-y

Wells, W. F. (1946). A Method for Obtaining Standard Suspensions of Tubercle Bacilli in the Form of Single Cells. Science, 104(2698), 254–255. https://doi.org/10.1126/science.104.2698.254

Wolf, A. J., Linas, B., Trevejo-Nunez, G. J., Kincaid, E., Tamura, T., Takatsu, K., & Ernst, J. D. (2007). Mycobacterium tuberculosis infects dendritic cells with high frequency and impairs their function in vivo. J Immunol, 179(4), 2509–2519. https://doi.org/10.4049/jimmunol.179.4.2509

